# Single-cell RNA transcriptome analysis reveals CXCL16/CXCR6 as maintenance factors for CNS tissue resident T cells that drive synapse elimination during viral recovery

**DOI:** 10.1101/2021.10.08.463715

**Authors:** Sarah F. Rosen, Allison L. Soung, Wei Yang, Shenjian Ai, Marlene Kanmogne, Veronica A. Davé, Sairam Prabhakar, Amanda Swain, Maxim Artyomov, Jeffrey A. Magee, Robyn S. Klein

**Affiliations:** Center for Neuroimmunology and Neuroinfectious Diseases, Washington University School of Medicine, St. Louis, MO; Department of Internal Medicine, Washington University School of Medicine, St. Louis, MO; Department of Genetics, Washington University School of Medicine, St. Louis, MO; Department of Pathology and Immunology, Washington University School of Medicine, St. Louis, MO; Department of Pediatrics, Washington University School of Medicine, St. Louis, MO; Department of Neuroscience, Washington University School of Medicine, St. Louis, MO

## Abstract

**Background:** Emerging RNA viruses that target the central nervous system (CNS) lead to cognitive sequelae in survivors. Studies in humans and mice infected with West Nile virus (WNV), a re-emerging RNA virus associated with learning and memory deficits, revealed microglial-mediated synapse elimination within the hippocampus. Moreover, CNS resident memory T (T_R_M) cells activate microglia, limiting synapse recovery and inducing spatial learning defects in WNV-recovered mice. The signals involved in T cell-microglia interactions are unknown.

**Methods:** Here, we examined the murine WNV-recovered forebrain using single-cell RNA sequencing to identify putative ligand-receptor pairs involved in intercellular communication between T cells and microglia. Clustering and differential gene analyses were followed by protein validation, genetic and antibody-based approaches utilizing an established murine model of WNV recovery in which microglia and complement promote ongoing hippocampal synaptic loss.

**Results:** Profiling of host transcriptome at 25 days post-infection revealed a shift in forebrain homeostatic microglia to activated subpopulations with transcriptional signatures that have previously been observed in studies of neurodegenerative diseases. Importantly, CXCL16/CXCR6, a chemokine signaling pathway involved in T_R_M cell biology, was identified as critically regulating CXCR6 expressing CD8^+^ T_R_M cell numbers within the WNV-recovered forebrain. We demonstrate that CXCL16 is highly expressed by all myeloid cells, and its unique receptor, CXCR6, is highly expressed on all CD8^+^ T cells. Using genetic and pharmacological approaches, we demonstrate that CXCL16/CXCR6 is required not only for the maintenance of WNV-specific, CD8 T_R_M cells in the post-infectious CNS, but also contributes to their expression of T_R_M cell markers. Moreover, CXCR6^+^CD8^+^ T cells are required for glial activation and ongoing synapse elimination.

**Conclusions:** We provide a comprehensive assessment of the role of CXCL16/CXCR6 as an interaction link between microglia and CD8^+^ T cells that maintains forebrain T_R_M cells, microglial and astrocyte activation, and ongoing synapse elimination in virally recovered animals. We also show that therapeutic targeting of CXCL16 during recovery may reduce CNS CD8^+^ T_R_M cells.

## Introduction

Neurotropic viruses may trigger the onset of neurodegenerative processes in the central nervous system (CNS) that lead to progressive memory impairments. West Nile virus (WNV) is an emerging neurotropic flavivirus that is the leading cause of domestically acquired arboviral disease and epidemic encephalitis in the United States^1^. Acute symptomatic syndromes include a self-limited febrile illness, West Nile fever (WNF), while more severe neuroinvasive diseases (WNND) include meningitis, encephalitis, or flaccid paralysis. Ninety percent of patients with WNND survive their acute illness due to the effector functions of antiviral T cells that clear virus in the CNS, predominantly via non-cytolytic cytokines, such as interferon (IFN)-γ^2–4^. The majority of these patients, however, develop debilitating cognitive and memory impairments that persist and worsen for years after recovery from encephalitis^5^. The mechanisms by which WNV infection in the CNS leads to long-term cognitive sequelae remain largely unknown.

Memory formation and consolidation is the result of complex circuitry between the hippocampus and cortex. The CA3 region of the dorsal hippocampus, in particular, plays an important role in the formation of spatial memories in rodents^6–8^. Using a novel murine model of WNV recovery, we previously traced spatial learning deficits to microglial engulfment of presynaptic terminals in the CA3 region of the hippocampus, acutely and ongoing, and is driven by classical complement protein C1qA expressed by neurons and microglia^9^. We also showed that CD8^+^IFNγ^+^ T cells that persist in the CNS as resident memory T (T_R_M) cells are the most proximal trigger of microglial activation, synapse elimination and spatial learning deficits during viral recovery^10^. The factors that determine which CD8^+^ T cells persist, how they interact with distinct CNS myeloid subsets, and whether this contributes to synapse elimination or cognitive impairment is not known.

To advance the discovery of cellular and molecular mechanisms that regulate differentiation and retention of CNS CD8^+^ T_R_M, we searched for genetic signatures of myeloid subpopulations and putative ligand-receptor regulators of microglia-T cell interactions, employing single-cell RNA sequencing (scRNA-seq) on forebrain cells (cortex and hippocampus) derived from WNV-recovered mice. Profiling of host transcriptome at 25 days post-infection revealed a shift in forebrain homeostatic microglia to activated subpopulations with transcriptional signatures similar to those previously observed in studies of neurodegenerative diseases^11–14^. Focusing on microglia and T cell subpopulations, selected genomic results underwent *in vivo* protein validation, followed by *in vivo* functional studies to define mechanisms that drive their interactions and functions.

Of the possible ligand-receptor pairs that drive intercellular interactions, cell localization and differentiation, chemokine signaling pathways are likely candidates. These molecules regulate the migration and recruitment of T cells during homeostasis and into inflamed tissues, where they also promote their differentiation^15, 16^. We identified *Cxcr6* as the only chemokine receptor in the top 20 differentially expressed genes (DEGs) within the CD8^+^ T cell cluster. CXCR6 is the unique receptor for the transmembrane chemokine CXCL16^17^; consistent with this, *Cxcl16* was highly expressed in all of the myeloid cell populations in the WNV-recovered CNS. CXCL16 is synthesized as a transmembrane multi-domain molecule, with a soluble version generated by cleavage of the transmembrane form through the actions of cell-surface proteases, such as a disintegrin and metalloproteinase (ADAM) 10 and 17^18–20^. CXCL16 is expressed in low levels by microglia during homeostatic conditions, and is highly expressed in the brain during pathological conditions such as multiple sclerosis, glioma, schwannomas and meningiomas^19, 21–25^. CXCR6 has been shown to play an important role in the localization and retention of T_R_M cells in the liver and lungs after viral infection^26–28^. Within the CNS, T_R_M cells that populate the human brain express CXCR6^29^, and meningeal γδ T cells have been shown to express high levels of CXCR6^30^. Using *Cxcr6^-/-^* mice and antibody-based neutralization of CXCL16, we demonstrate that CXCL16/CXCR6 are maintenance and differentiation factors for CD8^+^ T_R_M cells in the forebrain of WNV-recovered mice. Further, CXCR6 signaling promotes microglial activation and astrocytic interleukin (IL)-1β production, leading to ongoing synapse elimination in the CA3 region of the hippocampus. We provide novel genomic evidence for virus-mediated differentiation of neuropathologic microglial subsets, and the causal role of CXCL16/CXCR6 in hippocampal pathology during recovery from WNV CNS infection. We also provide evidence that targeting this pathway may be a successful approach for the prevention of microglial-mediated synapse loss that occurs after recovery from flavivirus encephalitis.

## Results

### scRNA-seq analysis identified eleven major cellular subtypes present in the WNV-recovered CNS

To investigate the responses of myeloid cell subpopulations and T cells after recovery from viral encephalitis and determine whether they share similar transcriptional signatures to those seen in neurodegenerative diseases in an unbiased manner, we performed scRNA-seq analyses on forebrain tissues (cortices and hippocampi) collected from C57BL/6 mock- and WNV-infected mice at 25 days post infection (DPI) (Fig. 1a). These studies used an established model of WNND, in which mice are infected intracranially with 10^4^ plaque-forming units (PFU) of an attenuated strain of WNV, WNV-NS5-E218A, which contains a single point mutation in the gene encoding 2′-O-methyltransferase, leading to type I interferon-mediated viral clearance by 15 DPI and 90% survival^31^. Forebrain tissue was collected for analysis as these brain regions are sites of processes that regulate learning and memory^6–9, 32^. A single-cell suspension was prepared from dissociated CNS tissue and transcriptomes of 4,250 cells were obtained. After data normalization, cells were clustered into eleven distinct clusters (Fig. 1b) using Seurat, and visualized using t-distributed stochastic neighbor embedding (t-SNE).

**Figure 1:**
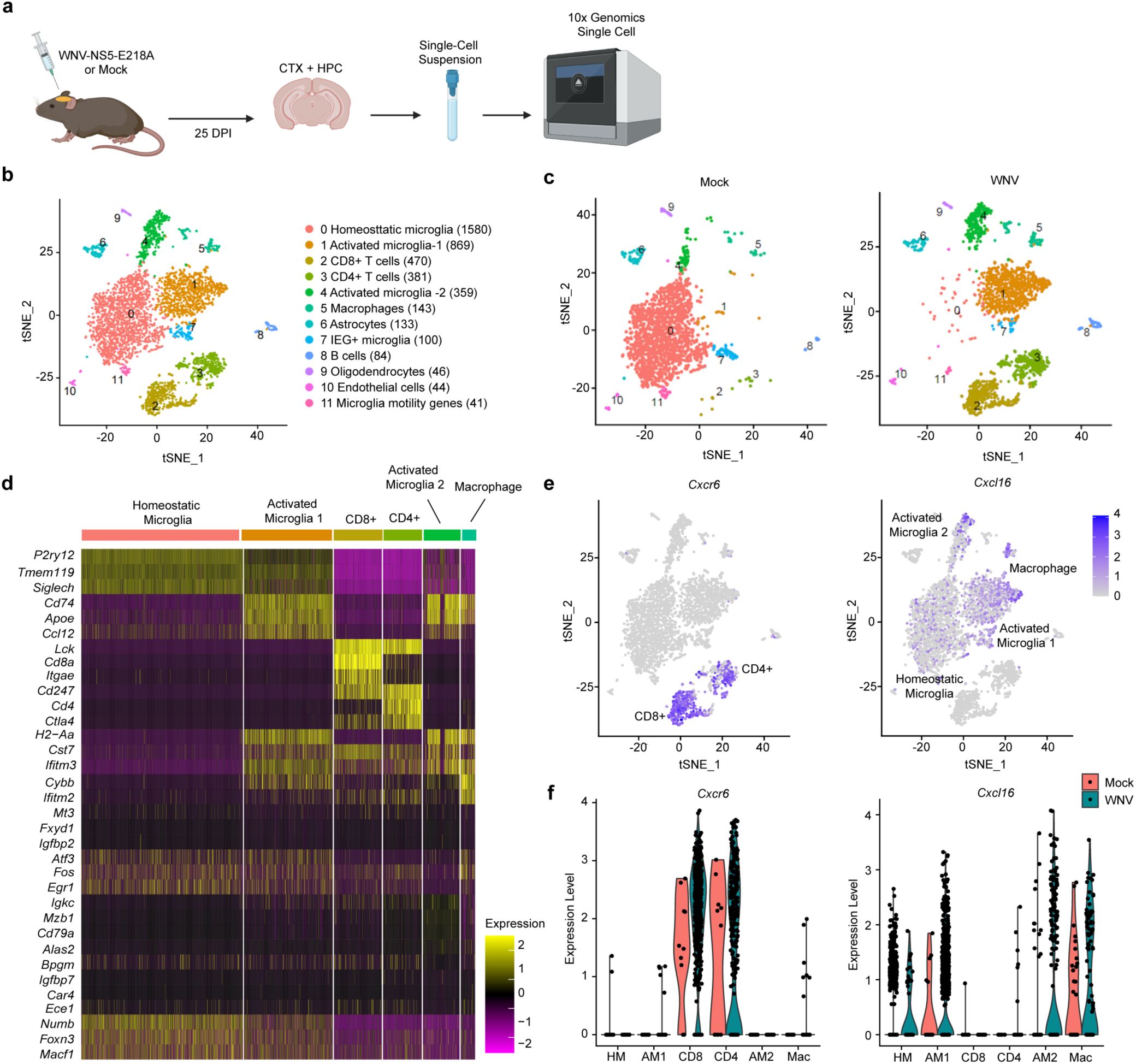
Clustering of CNS cells following WNV infection by scRNA-seq. **a.** Experimental design for scRNA-sequencing. Mice were infected (i.c.) with 1×10^4^ p.f.u. WNV-NS5-E218A and harvested 25 DPI. *N=*2 per group, each *N* includes 4 mice pooled. **b.** tSNE plot of all immune cells analyzed in both mock- and WNV-infected animals, showing eleven clusters, colored by density clustering, and annotated by cell-type identity and number of cells in parentheses. **c.** tSNE plots of all cells analyzed in mock (left panel)- and WNV (right panel)-infected mice, separated by treatment group, colored by density clustering. **d.** Heatmap of single cells representing the mRNA levels of the top three well-known genes used for cellular identification of each cluster. **e.** tSNE plot of all immune cells with mRNA of *Cxcr6* (left panel) and mRNA of *Cxcl16* (right panel). **f.** Violin plots showing immune cells with *Cxcr6* mRNA (left panel) or *Cxcl16* mRNA (right panel) in clusters 0-5. Each dot represents a cell.

Enriched genes for each cluster were determined by differential gene expression (DEG) analysis, and used to identify cell clusters. As expected, not all clusters were present in high numbers in both experimental groups (Fig. 1c). Major CNS cell-type identities for each cluster was assigned by cross-referencing these genes with published gene expression datasets^33–37^ (Fig. 1d, Supp. Table 1). The annotated eleven clusters included four microglia clusters, two T cell clusters, a macrophage cluster, an astrocyte cluster, and various other non-neuronal clusters. There were very low numbers of CD8^+^ (10) and CD4^+^ (11) T cells present in the mock-infected group. However, both CD8^+^ (460) and CD4^+^ (370) T cells were present in the WNV-infected group, consistent with previous data demonstrating very low levels of T cell entry into uninflamed CNS and T cell persistence in our model of WNV recovery^10, 38^. The homeostatic microglia cluster (Cluster 0) was predominantly present in CNS cells from mock-infected mice, expressing known markers of microglial homeostasis, including *P2ry12* (purinergic receptor P2Y12), *Siglech* (sialic acid binding Ig-like lectin H), and *Csfr1* (colony stimulating factor 1 receptor), while activated microglia clusters (Cluster 1 and 4) were enlarged in the CNS cells from the WNV-infected mice, expressing *Cd74* (cluster of differentiation 74), histocompatibility 2 class II antigen genes, *C4* (complement component 4), and *Apoe* (apolipoprotein E) (Table 1). Strikingly, the current dataset suggests that, even ten days after WNV is cleared from the CNS, almost all microglia in the WNV-infected mice remain in an activated state.

**Table 1.**
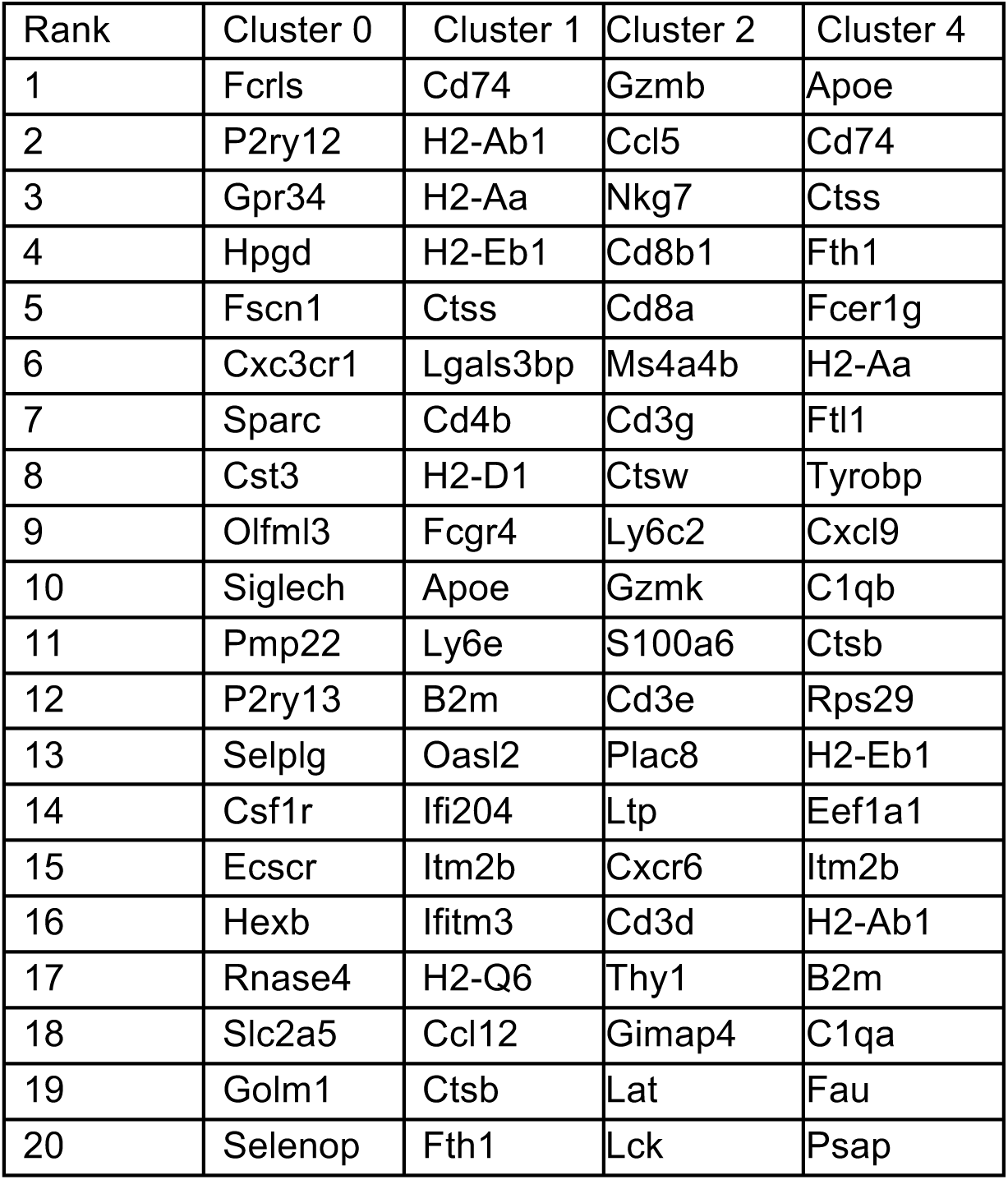
Top 20 defining genes of clusters 0,1,2, and 4.

### scRNA-seq analysis identified unique activated microglia clusters that persist in the forebrain of WNV-recovered mice

o investigate differences in the microglia clusters, we performed differential expression tests to compare microglia cluster 1 to 0, 4 to 0, and 4 to 1. A Venn diagram was created to show the number of significant genes from each of the 3 comparisons, and number of genes shared across comparisons (Supp. Fig 1a). There was a total of 842 differentially expressed genes from the comparison between cluster 1 and cluster 0, 482 genes between cluster 4 and 0, 319 between cluster 4 and 1. Of them, 272 genes were differentially expressing from both comparisons between 1 and 0 and between 4 and 0; 219 genes both from comparisons between 4 and 0 and between 4 and 1; 131 genes from both comparisons between 4 and 0 and between 1 and 0. 68 genes were differentially expressed across all 3 comparisons. Genes important for MHC and antigen presentation, such as *Cd74, H2-Ab1, H2-Eb1* were found among the top 25 significantly upregulated genes in cluster 1 when compared to cluster 0 (Supp. Fig. 1b) (Table 1). Homeostatic genes, such as *P2ry12, Cx3cr1, Fcrls, Siglech, and Hexb* were found among the top 25 significantly upregulated genes in cluster 0 compared to clusters 1 and 4 (Supp. Fig. 1c). Thus, both clusters 4 and 1 downregulate homeostatic microglial genes, but cluster 1 appears to upregulate more genes responsible for antigen responses than cluster 4. GO biological processes pathway analysis supported this, revealing upregulation in pathways involved in protein translation, antigen processing, and presentation of peptide antigen via MHC class I and class II in WNV cluster 1 vs mock cluster 0 and WNV cluster 4 vs mock cluster 0 (Supp. Fig. 2a,c). To further investigate the effects of WNV infection on microglia, we examined the levels of known homeostatic markers between cluster 0 and cluster 1 from both mock and WNV-infected mice^11, 39, 40^. *P2ry12, Cx3cr1, Tmem119, Fcrls, Siglech, Gpr34, and Hexb* were all significantly decreased in cluster 1 compared to cluster 0 (Supp. Fig. 2b), while cluster 4 was associated with upregulation of inflammatory molecules including *Ctss, Fth1, Tyrobp, Fcer1g, Cxcl9,* and *Apoe*, of which *Apoe* was one of the most upregulated genes (Table 1). These data reveal unique activated microglia signatures that persists in the CNS of WNV-recovered animals that exhibit similarities to those observed in studies using murine models of neurodegenerative diseases^40–42^.

### *Cxcl16* and *Cxcr6* are uniquely upregulated by microglia and T cells in the forebrain of WNV-recovered animals

Although T cells are critical for viral clearance from the CNS, their persistence in the CNS after viral clearance contribute to persistent microglial activation^32, 43^. Chemokine signaling pathways are important for regulating the recruitment and migration of leukocytes into the CNS, and during acute WNV infection infiltrating myeloid and T cells upregulate the expression of proinflammatory chemokine ligands and their receptors^44–47^. Thus, we manually sorted through DEGs in the myeloid and T cell clusters to investigate chemokine signaling pathways that are upregulated in the WNV-recovered forebrain. We identified *Cxcr6* as the only chemokine receptor in the top 20 DEGs within the CD8^+^ T cell cluster in the WNV-recovered forebrain (Table 1). *Cxcr6* was also highly upregulated in the CD4^+^ T cell cluster in the WNV-recovered CNS (Fig. 1e, f). CXCR6 is the unique receptor for the transmembrane chemokine CXCL16^17^. Consistent with this, *Cxcl16* was highly expressed in all of the myeloid cell populations in the WNV-recovered CNS (Fig 1e, f). These data suggest that the CXCL16/CXCR6 chemokine signaling axis may be important for T cell recruitment or maintenance within the CNS during WNV infection.

### CXCR6 is expressed by CD8^+^ T cells that persist in the CNS after viral recovery

To validate and extend our scRNA-seq findings, we performed an analysis of mock- versus WNV-infected CXCR6-EGFP (green fluorescent protein) reporter mice at 25 DPI. Results revealed a significant increase in CXCR6^+^ expressing cells in the forebrain of WNV-infected mice, which coincided with a significant increase in CXCR6^+^CD3^+^ co-localization via immunohistochemistry (Fig. 2a, b). In line with our scRNA-seq results, nearly 100% of the CXCR6^+^ expressing cells were shown to be CD3^+^, suggesting that T cells are the predominant source of CXCR6^+^ in the WNV-recovered CNS (Fig. 2a, b). Flow cytometric analysis of CD45^high^ cells isolated from the forebrain of wildtype (WT) WNV-infected mice at 7 (peak encephalitis), 25 (early recovery), and 52 (late recovery) DPI indeed revealed a significant increase in percentages of CD4^+^ and CD8^+^ T cells expressing CXCR6 at 25 and 52 DPI compared to 7 DPI (Fig. 2c, Supp. Fig. 3 a,c). While total numbers of CD4^+^ T cells expressing CXCR6 were still elevated at 25 DPI, significant differences in total numbers of CD8^+^ CXCR6^+^ T cells only 52 DPI (Supp. Fig 3a-d). Additionally, in the meninges at 52 DPI, approximately 100% of the CD8^+^ T cells are CXCR6^+^ (Supp. Fig. 4k). Further analysis revealed that approximately 100% of CD8^+^CXCR6^+^ in the CNS at 25 and 52 DPI express CD103, a marker for T_R_M cells (Fig. 2d), consistent with our scRNA-seq findings (Supp. Fig. 1d) and other studies linking CXCR6 to T_R_M cell biology^48^. Furthermore, 100% of CD8^+^IFNγ^+^ cells were CXCR6^+^ at 52 DPI (Supp Fig. 3e). MHC class I tetramer staining confirmed specificity of these CD8^+^ T cells to WNV NS4B, the immunodominant CD8 epitope in WNV-infected mice^49^, and approximately 100% of the CD8^+^NS4B^+^T cells were CXCR6^+^ in the CNS at 25 and 52 DPI (Fig. 2e).

**Figure 2:**
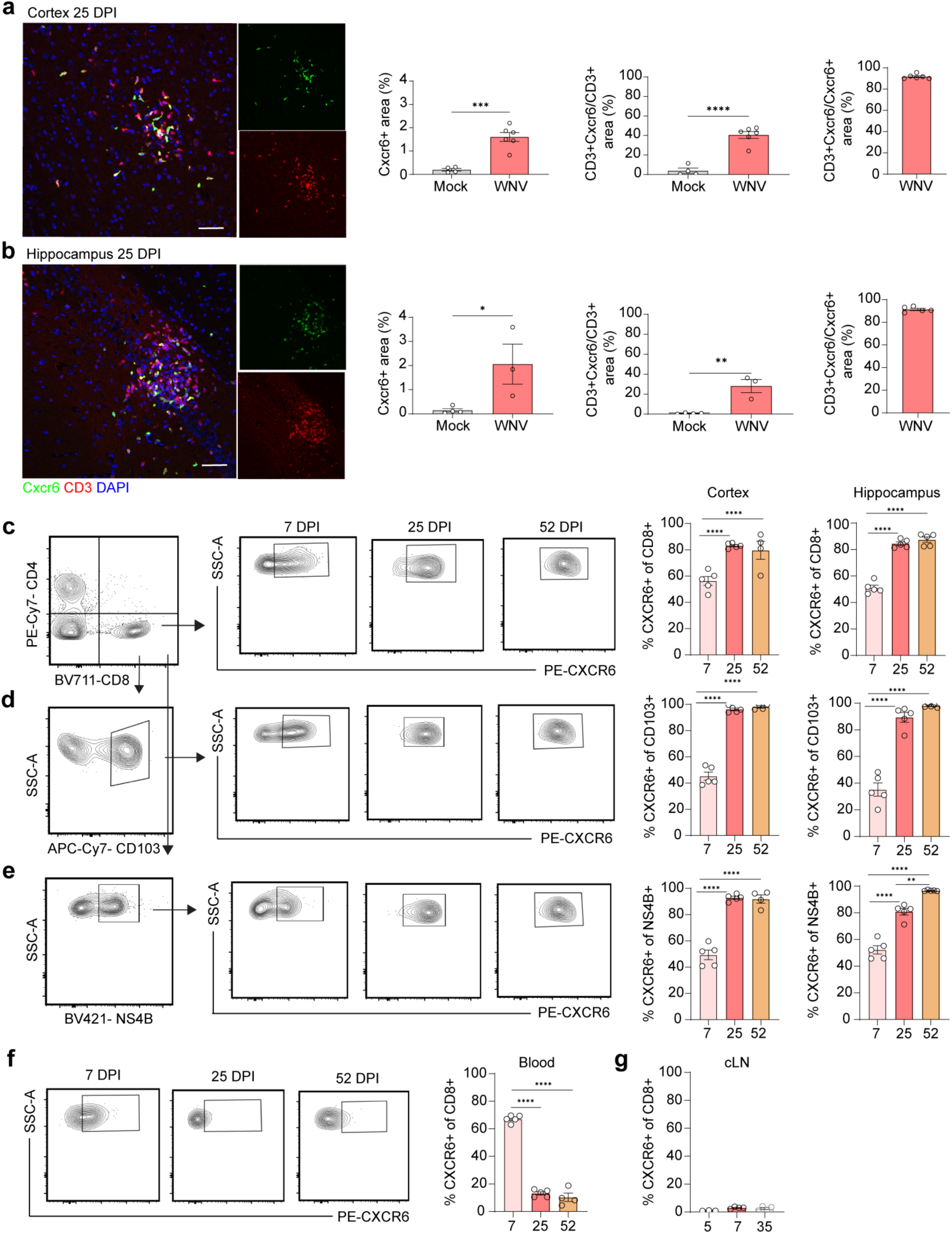
CXCR6 is expressed by CD8^+^ T cells that persist in the CNS after viral recovery. **a,b.** Immunohistological analysis for CXCR6 and CD3 in the cortex (a) and hippocampus (b) of mock- or WNV-infected mice at 25 DPI, presented as microscopy. Left graph shows percent CXCR6^+^ area, middle graph shows CD3^+^CXCR6^+^ area normalized to the total CD3^+^ area, indicative of co-localization, and right graph shows CD3^+^CXCR6^+^ area normalized to the total CXCR6^+^ area. Scale bars, 50 μm. **c.** Gating strategy and quantification of percentage of CD8^+^ T cells that are CXCR6^+^ at 7, 25 and 52 DPI in the cortex (left graph) and hippocampus (right graph). Cells were gated on CD45^high^ (Supp. Fig. 8a) before gating on CD4^+^ and CD8^+^ T cells (left panel). CD8^+^ T cells were then gated on CXCR6 (middle panels). **d.** Gating strategy and quantification of percentage of CD103^+^ cells that are CXCR6^+^ at 7, 25 and 52 DPI in the cortex and hippocampus. **e.** Gating strategy and quantification of percentage of NS4B^+^ cells that are CXCR6^+^ at 7, 25 and 52 DPI in the cortex and hippocampus. **f.** Gating strategy and quantification of percentage of CD8^+^ T cells that are CXCR6^+^ at 7, 25 and 52 DPI in the blood. Cells were gated on CD45^high^ before gating on CD4^+^ and CD8^+^ T cells. CD8^+^ T cells were then gated on CXCR6. **G.** Quantification of percentage of CD8^+^ T cells that are CXCR6^+^ at 7, 25 and 52 DPI in the cervical lymph nodes. Gating for CXCR6 based off *Cxcr6^-/-^* mice (Supp. Fig. 8b, c). Data represent the mean±s.e.m. and were analyzed by unpaired Student’s *t*-test. *P<0.05, **P<0.005, \*\*\**P* < 0.001, \*\*\*\**P* < 0.0001.

To determine whether CXCR6^+^ upregulation on CD8^+^ T cells occurs in the CNS or the periphery during WNV infection, we analyzed CXCR6 expression in the blood and cervical lymph nodes (cLN). CD8^+^ T cells in the blood upregulate CXCR6 at 7 DPI, with about 65% of CD8^+^ T cells expressing CXCR6. However, during recovery at 25 and 52 DPI, only 8% of the CD8^+^ T cells in the blood express CXCR6 (Fig. 2f). The percentage of CD8^+^ CXCR6^+^ T cells remained at about 5% in the cLN, and did not increase significantly throughout the duration of viral recovery. Of note, CD4^+^ CXCR6^+^ T cell percentages or numbers were not increased in the blood at any time point (Supp. Fig. 3f). Taken altogether, these data suggest that high percentages of CD8^+^ CXCR6^+^ T cells are found within the blood as the cells traffic into the CNS via a route that includes the meninges, and that they persist in the CNS throughout recovery.

### *Cxcl16* levels increase on IBA1^+^ cells during acute infection and return to baseline levels during recovery

To validate scRNA-seq detection of *Cxcl16* mRNA expression within the WNV-recovered CNS, we performed *in situ* and immunohistochemical analysis of CNS tissues derived from mock- and WNV-infected mice at 7, 25, and 52 DPI. There was a significant increase in *Cxcl16* mRNA in WNV-infected tissue at 7 and 25 DPI compared to mock (Fig. 3a, b). Co-localization of *Cxcl16* within IBA1^+^ cells detected increased levels in the cortex at 7 and 52 DPI, and the hippocampus at 7 and 25 DPI, compared to mock (Fig. 3c). Within the CNS, nearly 100% of the *Cxcl16* is co-localized with IBA1 at 7, 25, and 52 DPI, suggesting that myeloid cells are the predominant cellular source of *Cxcl16*. (Fig. 3c, d). Notably, *Cxcl16* mRNA is found in the soma and processes of IBA1^+^ cells at 7 DPI (Fig. 3a, white box inset), suggesting local translation of *Cxcl16* within cellular processes. These data indicate that *Cxcl16* is expressed by IBA1^+^ cells within the CNS. To determine whether CXCL16 protein was being produced in the parenchyma of the CNS, we performed ELISA on cortical and hippocampal tissue, meninges, blood and lymph nodes in mock and WNV-infected mice at 7 and 25 DPI. CXCL16 levels were increased at 7 and 25 DPI in the cortex and 7 DPI in the hippocampus compared to mock. However, there was also a modest increase in CXCL16 in the cervical lymph nodes at 25 DPI (Fig. 3e), indicative of a peripheral source of CXCL16 during recovery that may be competing with CNS sources of CXCL16 to recruit CD8^+^ T cells out of the CNS. To determine whether myeloid cells were interacting with CXCR6^+^ T cells within the CNS, we examined CXCR6-EGFP reporter mice at 25 DPI, and observed CD3^+^CXCR6^+^ T cells in close proximity to IBA1^+^ cells (Fig. 3f). Together, these data suggest that CXCL16 from IBA1^+^ myeloid cells may direct interactions with CD3^+^ CXCR6^+^ cells during CNS infection and recovery.

**Figure 3.**
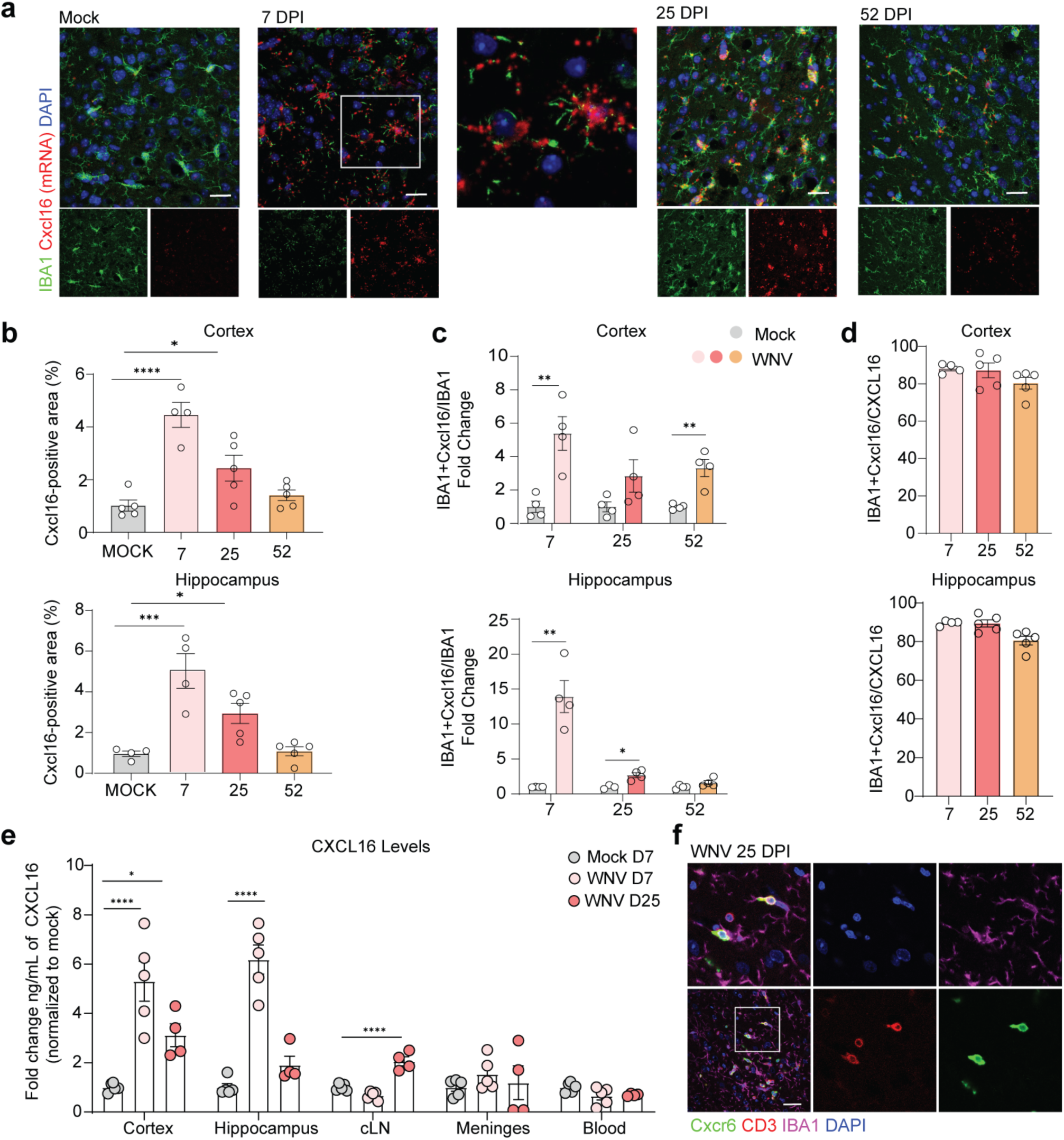
CXCL16 levels increase on IBA1^+^ cells during acute infection and return to baseline levels during recovery. **a.** Representative microscopy images of RNA *in situ*/immunohistochemistry (ISH/IHC) for *Cxcl16* and IBA1 in the cortex of mock- or WNV-infected mice at 7, 25 ad 52 DPI. Middle panel is inset of white box in 7 DPI. **b-d.** Quantification of ISH/IHC for *Cxcl16^+^* area (b), IBA1^+^*Cxcl16^+^* area normalized to the total IBA1^+^ area, represented as fold change over mock (c), or IBA1*^+^Cxcl16^+^* area normalized to the total *Cxcl16*^+^ area (d) in the cortex or hippocampus of mock- or WNV-infected mice at 7, 25 and 52 DPI. **e.** ELISA for CXCL16 in the cortex, hippocampus, cervical lymph nodes, meninges, and blood at 7 and 25 DPI. **f.** Representative immunostaining of the cortex for CXCR6-GFP, CD3, IBA1, and DAPI. Scale bars, 50 μm. Data represent the mean±s.e.m. and were analyzed by one-way ANOVA or unpaired Student’s t-test. *P<0.05, **P<0.005, ***P < 0.001, ****P < 0.0001.

### CXCR6 is not required for CNS virologic control during acute WNV infection

To determine whether CXCR6 contributes to antiviral T cell responses within the CNS during acute infection, we examined immune cell infiltration and viral clearance in WT and *Cxcr6^-/-^* mice. First, we confirmed the deletion of CXCR6 on both CD8^+^ and CD4^+^ T cells in the forebrain of *Cxcr6^-/-^* mice (Supp. Fig. 4e, f). Analysis of CD45^high^ cells at 7 DPI revealed no differences in frequency or number of CD4^+^, CD8^+^, CD103^+^, or NS4B^+^ T cells in the forebrain between WT and *Cxcr6^-/-^* mice (Fig. 4a, b, c. Supp. Fig. 4h,i). There were also no significant differences in the percentages of CD8^+^CXCR6^+^ or CD4^+^CXCR6^+^ T cells in the blood between WT and *Cxcr6^-/-^* mice at 7 DPI (Supp. Fig. 4j). Furthermore, WNV infection of WT and *Cxcr6^-/-^* mice did not reveal significant differences in survival, weight loss, acute levels or clearance of viral loads in the CNS (Supp. Fig. 4a, b, c, d). These data suggest that CXCR6 is not necessary for the trafficking and antiviral functions of T cells in the CNS.

**Figure 4.**
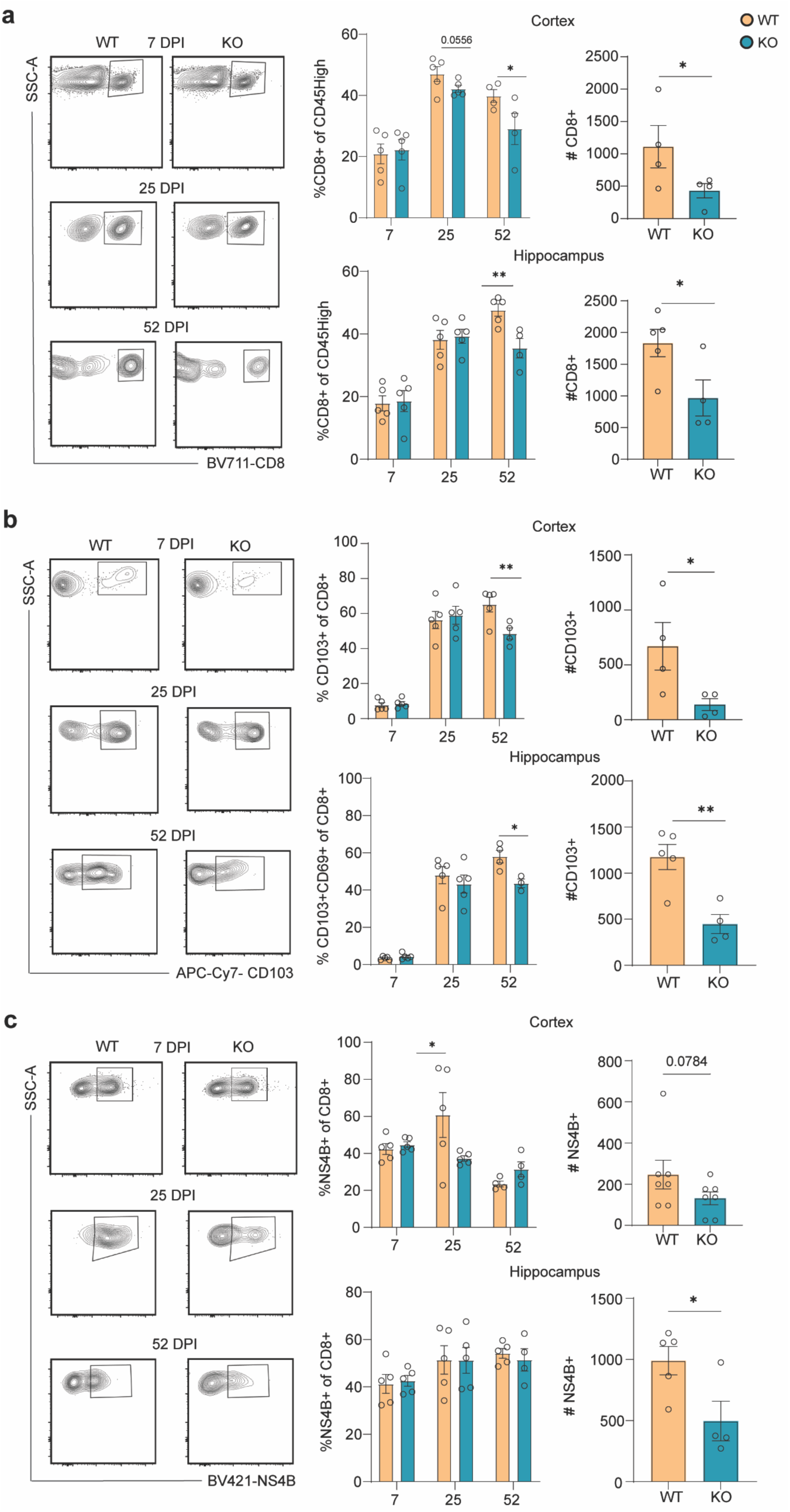
*Cxcr6^-/-^* animals have decreased maintenance of CD8^+^ T cells in the post-infectious CNS. **a.** Representative gating strategy and quantification of percentage of CD8^+^ T cells within the cortex and hippocampus at 7, 25 and 52 DPI (left graph), followed by total CD8^+^ cell numbers at 52 DPI in WT versus *Cxcr6^-/-^* mice (right graph). **b.** Representative gating strategy and quantification of percentage of CD103^+^ cells within the cortex and hippocampus at 7, 25 and 52 DPI, followed by total CD103^+^ cell numbers at 52 DPI, in WT versus *Cxcr6^-/-^* mice. **c.** Representative gating strategy and quantification of percentage of NS4B^+^ cells within the cortex and hippocampus at 7, 25 and 52 DPI, followed by total NS4B^+^ cell numbers at 52 DPI in WT versus *Cxcr6^-/-^* mice. Cells were gated according to gating strategy in Supp. Fig. 8. Data represent the mean±s.e.m. and were analyzed by unpaired Student’s *t*-test at 7, 25 and 52 DPI. *P<0.05, **P<0.005.

### CXCR6 is required for the maintenance of CD8^+^ T_R_M cells in the forebrain after recovery from WNV

Given the high expression of CXCR6 on T cells within the CNS after viral recovery, we hypothesized that CXCR6 may play an important role in maintenance of CD4^+^ and/or CD8^+^ T cells in the CNS. To test this hypothesis, we performed phenotypic analyses of infiltrating T cells in the CNS and peripheral tissue from mock- and WNV-infected wild-type (WT) and *Cxcr6^-/-^* mice at 25 and 52 DPI. At 25 DPI, there was a trend towards a decrease in percentage of CD8^+^ T cells and a significant decrease in the percentage of CD8^+^NS4B^+^ T cells in the cortex in *Cxcr6^-/-^* compared with WT animals (Fig. 4a, d). However, there were no differences in percentage of CD103^+^ or numbers of CD8^+^, CD103^+^, or NS4B^+^ in the cortex at 25 DPI (Fig. 4a, b, c, Supp. Fig. 5b). Similarly, there were no significant differences in percentages or numbers of CD8^+^, CD103^+^, or NS4B^+^ in the hippocampus at 25 DPI (Fig. 4a, b, c, Supp. Fig. 5b). In contrast, at 52 DPI, there were significant decreases in both percentages and numbers of CD8^+^ and CD8^+^CD103^+^ cells in the both CNS regions of WNV-recovered *Cxcr6^-/-^* mice compared with WT mice (Fig. 4a, b). Of note, the percentages of CD8^+^NS4B^+^ cells were not significantly different at 52 DPI in either brain region, nor were there differences in percentages of CD103^+^CD8^+^ that were NS4B^+^ at 7, 25, or 52 DPI, suggesting that CXCR6 is not required for the maintenance of virus-specific CD8^+^ T cells (Fig. 4c, Supp. Fig. 4g). However, the numbers of CD8^+^NS4B^+^ T cells were significantly lower in both the hippocampi and cortices of *Cxcr6^-/-^* mice compared to WT mice at 52 DPI, which is likely a result of a decrease in total CD8^+^ T cells in the forebrain (Fig. 4c). We also did not see any significant differences in CD4^+^ T cell numbers in the CNS at 25 or 52 DPI between WT and *Cxcr6^-/-^* mice (Supp. Fig. 4h, i). Analysis of the blood did not reveal any significant differences in the percentages or numbers of CD8^+^CXCR6^+^ or CD4^+^CXCR6^+^ cells between WT and *Cxcr6^-/-^* mice at 25 or 52 DPI (Supp Fig. 4j). These data suggest that CXCR6 is necessary for the persistence of CD8^+^ T cells in the CNS after recovery from viral infection.

Terminal deoxynucleotidyl transferase dUTP nick end labeling (TUNEL) and CD3^+^ immunodetection within the hippocampi and WNV-recovered WT and *Cxcr6^-/-^* mice at 52 DPI did not reveal any differences between the genotypes (Supp. Fig. 6a). This suggests that increased rates of apoptosis are not responsible for the loss of T_R_M cells in the CNS of *Cxcr6^-/-^* mice. Similarly, analysis of hippocampi from WT and *Cxcr6^-/-^* CD8^+^ T cell at 35 DPI for expression of CCR7 and CD62L, a chemokine receptor and selectin, respectively, with known roles in facilitating the homing of memory CD8^+^ T cells back to lymph nodes^50, 51^, did not reveal any significant differences between the genotypes (Supp. Fig. 6b). For these studies, day 35 was chosen as a midpoint between 25 to 52 DPI, the latter of which was when we detected significantly decreased percentages and numbers of CD8^+^ T cells in the forebrain of *Cxcr6^-/-^* mice. Finally, studies quantitating CD8^+^ T cells within the meninges of WNV-recovered WT and *Cxcr6^-/-^* mice did not indicate that the decreased CD8^+^ T cells in the forebrain of *Cxcr6^-/-^* mice was due to their accumulation within the meninges (Supp. Fig. 4l). Together, these results suggest that the loss of CD8^+^ T cells from the CNS of *Cxcr6^-/-^* mice is not attributable to substantial differences in apoptosis, egress to the lymph node, or accumulation in the meninges.

### CXCR6 signaling contributes to persistent microglial activation and impaired synapse recovery in the hippocampi of WNV-infected animals

We previously showed that CD8^+^ T_R_M cell-derived IFNγ signaling is required for microglial activation and astrocyte expression of interleukin(IL)-1β within the hippocampus of WNV-recovered mice^10^. Given the significant decrease in CD8^+^CD103^+^ cells in the hippocampi of WNV-recovered *Cxcr6^-/-^* mice, we hypothesized that CXCR6-deficient mice would exhibit attenuation of these responses during recovery from WNV infection. Consistent with prior studies^52^, we detected increased IL-1β expression by GFAP^+^ astrocytes in the hippocampi of WNV-infected WT mice at 52 and 25 DPI compared with mock-infected animals (Fig. 5a, Supp. Fig. 7a, respectively). Although astrocyte expression of IL-1β was also elevated in *Cxcr6^-/-^* mice at 25 DPI compared with mock-infected animals, it was significantly lower than that observed in WNV-infected WT mice at 52 DPI (Fig. 5a). Similarly, while WNV infection led to increased IBA1^+^ expression within the hippocampus of WT and *Cxcr6^-/-^* mice at 25 DPI compared to mock (Supp. Fig. 7a), at 52 DPI *Cxcr6^-/-^* mice exhibited significantly less IBA1^+^ expression the hippocampus compared to similarly infected WT mice (Fig. 5b). These data are consistent with the findings that WNV-specific CD8^+^ T cells are found in similar numbers in the CNS of WT and *Cxcr6^-/-^* mice at 25 DPI, but are significantly lower in *Cxcr6^-/-^* mice at 52 DPI (Fig. 4a). Notably, while WNV-recovered *Cxcr6^-/-^* mice exhibited significantly higher IBA1^+^ expression compared to mock-infected controls, there were no differences in detection of phagocytic IBA1^+^CD68^+^ cells. In contrast, WNV-recovered WT mice exhibited significant increases in IBA1^+^CD68^+^cells compared to WT mock and WNV-recovered *Cxcr6^-/-^* mice (Fig. 5b). Together, these data suggest that CD8^+^CXCR6^+^ T cells in the hippocampus play a role in maintaining phagocytic microglia and astrocyte expression of IL-1β within the hippocampus of WNV-infected animals during recovery.

**Figure 5.**
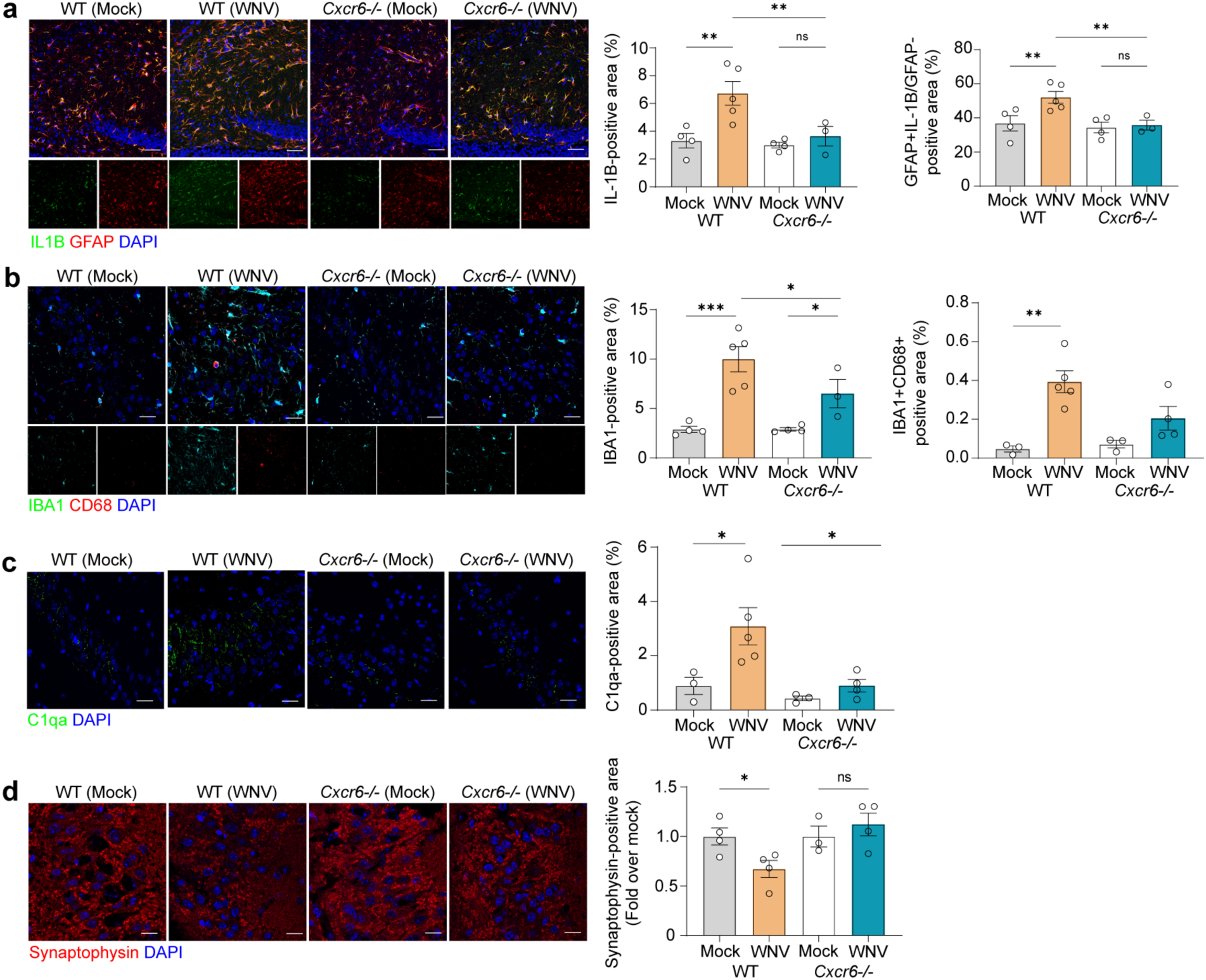
CXCR6 signaling potentiates gliosis in the recovered hippocampi of WNV-infected animals, contributing to synaptic elimination. **a.** Representative immunostaining and quantification of IL-1β (green), GFAP (red) and DAPI (blue) in CA3 region of the hippocampus of mock or WNV-infected WT or *Cxcr6^-/-^* mice at 52 DPI. GFAP quantified by percent positive area, followed by quantification of GFAP^+^IL-1β^+^ area, normalized to the total GFAP^+^ area. **b.** Representative immunostaining and quantification of IBA1 (green), CD68 (red), and DAPI (blue) in hippocampus of mock- or WNV-infected WT or *Cxcr6^-/-^* animals at 52 DPI. IBA1^+^ quantified as percent positive area, followed by quantification of IBA^+^CD68^+^ percent positive area. **c.** Representative immunostaining and quantification of C1qa in the hippocampus of mock- or WNV-infected WT or *Cxcr6^-/-^* animals at 52 DPI showing staining for C1qa (green) and DAPI (blue). C1qa quan- tified as percent positive area. **d.** Representative immunostaining and quantification of synapses in the CA3 region of the hippocampus in mock- or WNV-infected WT or *Cxcr6^-/-^* animals at 52 DPI showing staining for synaptophysin (red) and DAPI (blue). Synaptophysin quantified by percent positive area. Scale bars, 50 μm (a-c) or 20 μm (d). Data represent the mean±s.e.m. and were analyzed by two-way ANOVA and corrected for multiple comparisons. *P<0.05, **P<0.005, \*\*\**P* < 0.001.

Given the significant decrease in IBA1^+^CD68^+^ cells within the hippocampi of WNV-recovered *Cxcr6^-/-^* mice compared to WT animals, we wondered whether CXCR6-deficient mice would also exhibit decreased complement and microglial-mediated synapse elimination, which was especially profound in the CA3 region of hippocampi in WT animals^9^. At 25 DPI, both WNV-recovered WT and *Cxcr6^-/-^* mice exhibited loss of presynaptic terminals, as assessed by detection of synaptophysin (Supp. Fig. 7c). However, at 52 DPI only WT mice continued to exhibit decreased presynaptic terminals compared to mock (Fig. 5d). Additionally, at 52 DPI, WNV-recovered *Cxcr6^-/-^* mice exhibited similar levels of synaptophysin as mock-infected *Cxcr6^-/-^* mice, and immunohistochemical examination of C1QA protein within the CA3 region of the hippocampus in mock-versus WNV-recovered mice revealed no differences in *Cxcr6^-/-^* mice, but a significant increase in WT WNV-recovered mice (Fig. 5c). Together, these data suggest that decreases in C1QA, microglial activation and astrocyte IL-1β expression prevented ongoing synapse elimination (Fig. 5d, Supp. Fig. 7c). These studies demonstrate a critical role for CXCR6 in promoting injury to the hippocampal synaptic circuit through the regulation of CD8^+^ T cell number and microglial activation within the WNV-recovered hippocampus.

### CD8^+^CXCR6^+^ T cells maintain T_R_M cells and synapse elimination in the forebrain after recovery from WNV infection

To confirm that CXCR6 specifically on CD8^+^ T cells is required for maintenance of T_R_M cells, microglial activation, and synapse elimination in the hippocampus after WNV recovery, naïve CD8^+^ T cells were isolated from the spleens of WT or *Cxcr6^-/-^* mice and adoptively transferred (AT) into *Cd8^-/-^* mice. *Cd8^-/-^* mice were infected 24 hours after AT, followed by harvesting of CD8^+^ T cells from the CNS at 52 DPI (Fig. 6a). Phenotypic analysis revealed that recipients of CD8^+^ T cells derived from *Cxcr6^-/-^* mice exhibited significantly lower percentages of CD8^+^NS4B^+^, and CD8^+^CD103^+^ cells in the forebrain than those that received WT CD8^+^ T cells (Fig. 6b, c). Upon examination of the CD8^+^ T cells that did persist within the CNS of mice that received WT CD8^+^ T cells, CXCR6 was expressed on approximately 100% (Fig. 6b), confirming that CXCR6 is essential for the generation of T_R_M cells. Importantly, mice that received CD8^+^ T cells from *Cxcr6^-/-^* mice also exhibited significantly lower immunodetection of IBA1^+^ and IBA1^+^CD68^+^ percent area compared to mice that received WT CD8^+^ T cells (Fig. 6d). Finally, mice that received *Cxcr6^-/-^* CD8^+^ T cells exhibited significantly higher levels of synaptophysin than those that received WT CD8^+^ T cells (Fig 6e). Together, these data suggest a cell-intrinsic role for CXCR6 in promoting CD8^+^ T_R_M cell maintenance in the CNS after viral encephalitis, contributing to persistent microglial activation and synaptic elimination.

**Figure 6.**
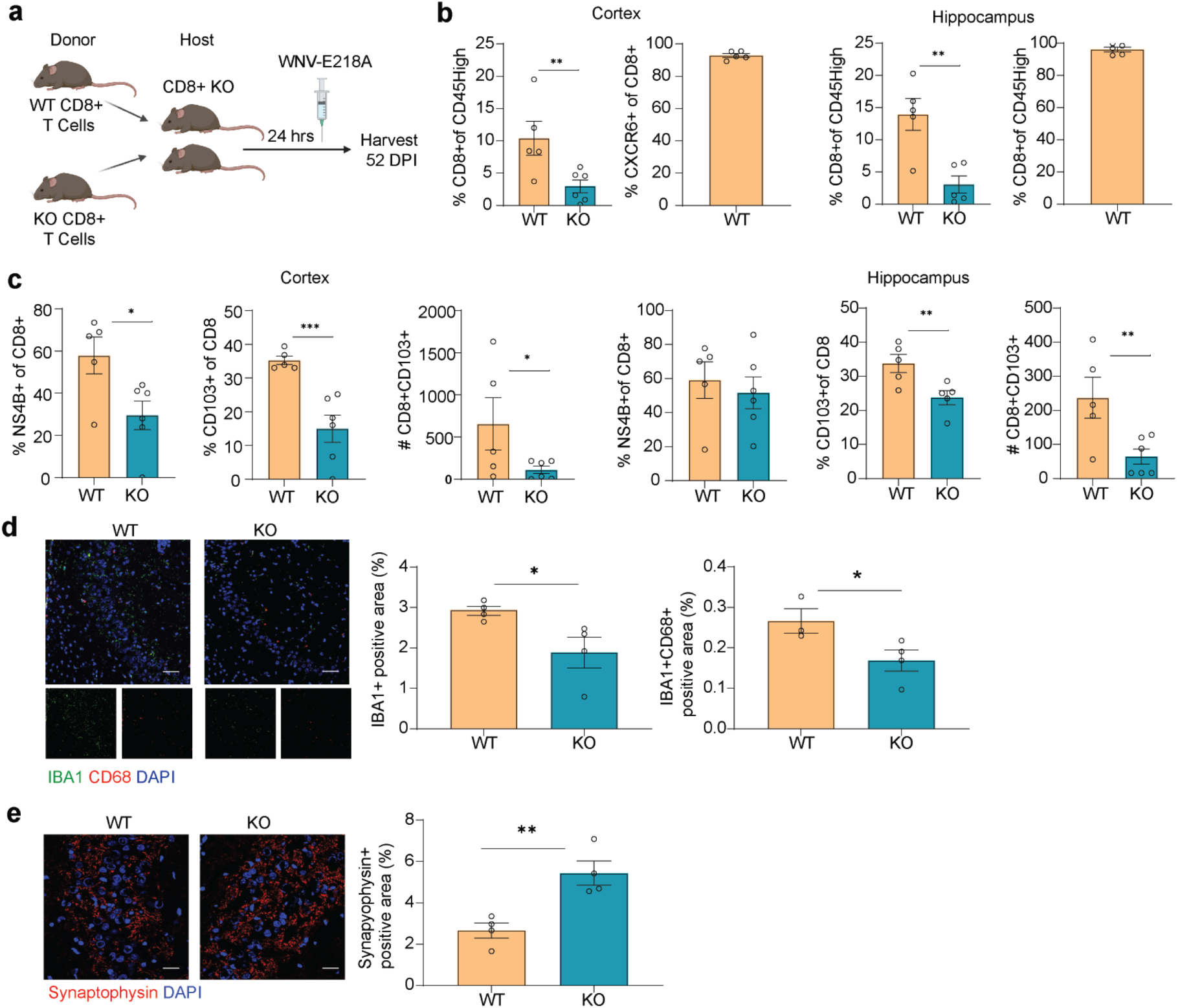
CXCR6 on CD8^+^ T cells is necessary for the maintenance of T_R_M cells in the CNS after recovery from WNV infection. **a.** Schematic showing experimental design of CD8^+^ T cell adoptive transfer from WT and *Cxcr6^-/-^* mice into *Cd8^-/-^* mice. 24 hours after adoptive transfer, mice were infected (i.c.) with 1×10^4^ p.f.u. WNV-NS5-E218A and harvested at 52 DPI. **b.** Flow cytometric analysis of the cortex and hippocampus at 52 DPI quantifying the percentage of CD45^high^ cells that are CD8^+^ in WNV-infected mice that received WT or *Cxcr6^-/-^* CD8^+^ T cells, followed by the percentage of CD8^+^ T cells that are CXCR6^+^ at 52 DPI in the mice that received WT CD8^+^ T cells. **b.** Flow cytometric analysis of the hippocampus and cortex at 52 DPI quantifying the percentage of CD8^+^ T cells that are NS4B^+^ or CD103^+^ in WNV-infected mice that received WT or *Cxcr6^-/-^* CD8^+^ T cells, followed by the total number of CD8^+^CD103^+^ T cells at 52 DPI in WNV-infected mice that received WT or *Cxcr6^-/-^* CD8^+^ T cells. **d.** Representative immunostaining and quantification at 52 DPI of IBA1 (green), CD68 (red), and DAPI (blue) in the hippocampus of WNV-infected mice that received WT or *Cxcr6^-/-^* CD8^+^ T cells. IBA1^+^ quantified as percent positive area, followed by quantification of IBA^+^CD68^+^ percent positive area. **e.** Representative immunostaining and quantification at 52 DPI of synapses in the CA3 region of the hippocampus of WNV-infected mice that received WT or *Cxcr6^-/-^* CD8^+^ T cells showing staining for synaptophysin (red) and DAPI (blue). Synaptophysin quantified by percent positive area. Scale bars, 50 μm. Data represent the mean±s.e.m. and were analyzed by unpaired Student’s *t*-test. *P<0.05, **P<0.005, ***P < 0.001.

### CXCL16 neutralization leads to decreased frequency of T_R_M cells in the WNV-recovered forebrain

Given our findings that CXCR6 signaling generates and maintains CD8^+^CD103^+^ T cells in the forebrains of WNV-recovered, and that this is required for neuropathology that contributes to cognitive dysfunction, we hypothesized that exogenous neutralization of CXCL16 might eliminate CD8^+^ T_R_M cells. To test this hypothesis, we intravenously administered CXCL16 neutralizing monoclonal antibody or isotype control to WT WNV-infected mice at 7, 8, 9, and 10 DPI (Fig. 7a). These time points were chosen based on published data indicating that permeability of the blood-brain barrier peaks at 9 DPI in our model of WNV recovery^52^. CNS tissues were harvested at 15 DPI, a time-point when differentiating T_R_M cells responding to acute infections robustly express CD103^53^. Phenotypic analysis of forebrain T cells at 15 DPI demonstrated significantly lower percentage of CD8^+^CD103^+^ T cells in the cortex of the anti-CXCL16 antibody-treated group, and a similar trend in the hippocampus (Fig. 7b, c). As expected, there were no significant decreases in the percentages of total CD8^+^ T cells in the forebrain of the anti-CXCL16 antibody-treated group. These data indicate that antibody-based neutralization of CXCL16 limits the generation of fore-brain CD8^+^CD103^+^ T cells and suggests CXCL16 might be a therapeutic target.

**Figure 7.**
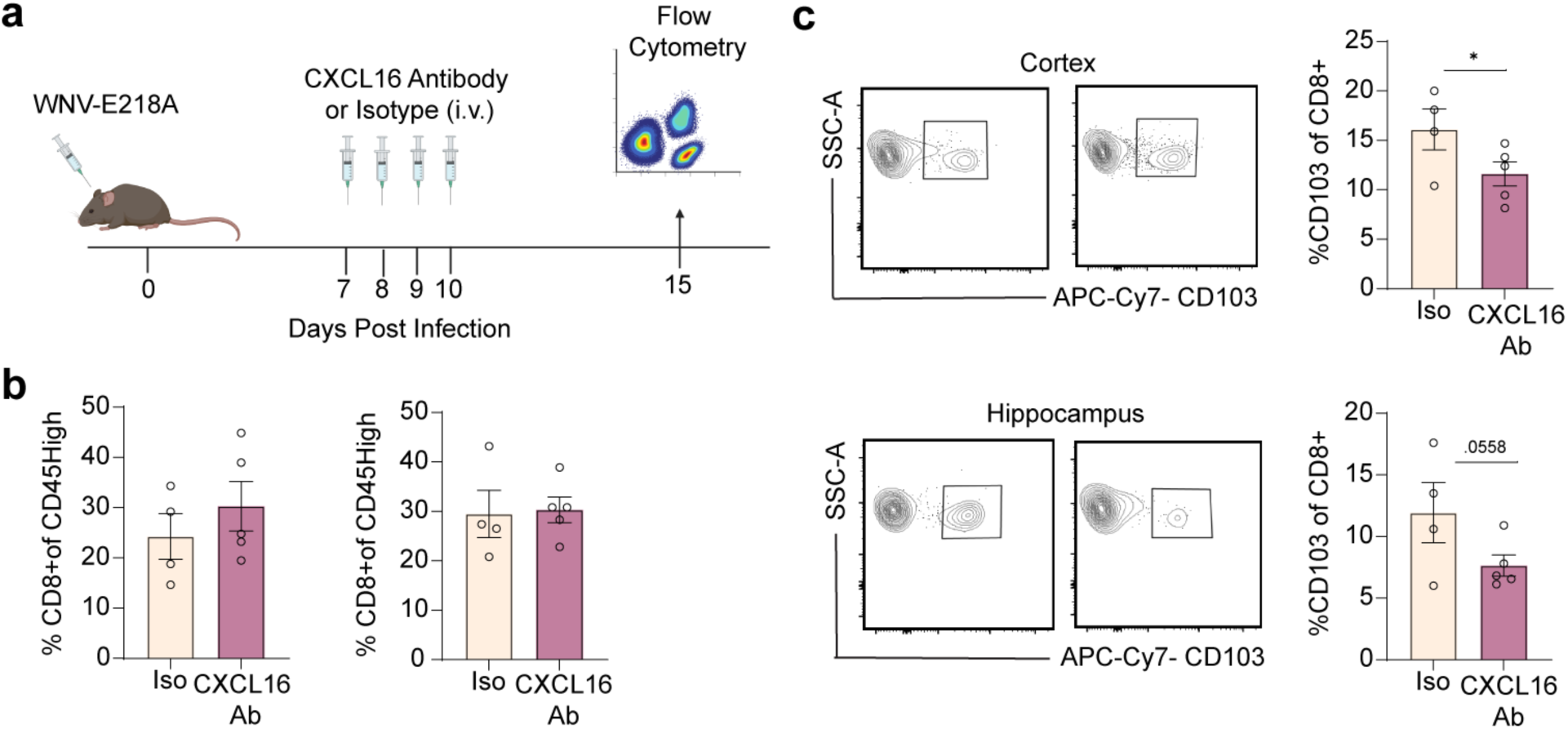
CXCL16 neutralization leads to decreased percentage of T_R_M cells in the CNS after viral clearance. **a.** Schematic depicting experimental design for CXCL16 neutralizing antibody experiment. Mice were infected (i.c.) with 1×10^4^ p.f.u. WNV-NS5-E218A and administered CXCL16 antibody or isotype control via retro-orbital injection, and harvested 15 DPI. **b.** Quantification of percentage of CD45^high^ cells that are CD8^+^ in the cortex (left) and hippocampus (right) in mice that received isotype or CXCL16 neutralizing antibody. **c.** Gating strategy and quantification of percentage of CD8^+^ T cells that are CD103^+^ in the cortex and hippocampus in mice that received isotype or CXCL16 neutralizing antibody. Data represent the mean±s.e.m. and were analyzed by unpaired Student’s *t*-test. *P<0.05.

## Discussion

During viral encephalitis, infection of neural cells leads to the activation of resident glial cells, such as microglia and astrocytes, induction of cytokines and chemokines, as well as leukocyte invasion into the CNS^8^,. The extent to which acute infection and inflammation contribute to disease pathology during recovery is complex, involving the coordination of multiple cell types and their sub-populations, each with distinct functions^58–60^. Prior to the advent of scRNA-seq, which enables profiling of single cells without prior gene knowledge and clustering based on transcriptional signatures, unbiased investigation of cells that persist within the CNS after recovery from viral encephalitis, and their functional consequences, has been difficult. Here, we identified eleven cell clusters present in the WNV-recovered forebrain, including four microglia clusters, two T cell clusters, a macrophage cluster, an astrocyte cluster, and various other non-neuronal clusters. It is important to note that relatively fragile cells, such as neurons and oligodendrocytes, are often reduced or lost during processing for scRNA-seq^58^. Thus, the 11 cell clusters identified here are not a complete representation of all of the cell types present in the WNV-recovered forebrain. Of the myeloid clusters identified, we detected unique activated microglia clusters that express the chemokine *Cxcl16*, and found that all persisting CD8^+^ T cells and many CD4^+^ T cells express CXCL16’s only receptor, CXCR6. Using mice with global deletion of *Cxcr6* or intravenous administration of CXCL16 neutralizing antibodies, we showed that CXCL16/CXCR6 differentiate and maintain forebrain T_R_M cells, and are required for microglial and astrocyte activation, and ongoing synapse elimination in virally recovered animals

Microglia are CNS-resident mononuclear phagocytic cells that are important for both immune protection via expression of pattern recognition receptors (PRRs), and neuronal function by regulating homeostasis and synaptic remodeling^59^. ScRNA-seq analyses of microglia has previously demonstrated their heterogeneity, and that they adapt their gene expression profiles during development, homeostasis, and perturbations^11, 60^. Microglia in the uninflamed brain exhibit a homeostatic genetic signature, expressing mRNAs for *P2ry12, Cx3cr1, Temem119, Siglech, Hexb,* and *Fcrls,* which are all downregulated during aging and in neurodegenerative diseases associated with cognitive disruption^61, 62^. We see that recovery from CNS infection with WNV, which may induce spatial learning defects, is also associated with down-regulation of these homeostatic microglial gene signatures. Consistent with this this, activated microglia cluster 1 instead upregulates genes involved in antiviral defense, peptide metabolic processes, antigen processing and presentation, and protein synthesis, such as *Cd74, Ifitm3, H2-Ab1, B2m, Ctss,* and *Apoe,* while cluster 4 additionally expressed high levels of *Tyrobp, Fcer1g, and Cxcl9*. Notably, microglial expression of these genes has been implicated in aging or in neurodegenerative disorders such as Alzheimer’s and Parkinson’s diseases, and Multiple Sclerosis^40–42, 63, 64^. These data suggest viral encephalitis may be a trigger for progressive neurodegenerative diseases.

Upon investigation of the top 20 genes in the T cell clusters, we identified *Cxcr6* as one of the top DEGs in the CD8^+^ T cell cluster. CXCR6 is the unique receptor for CXCL16, which is upregulated in activated microglia clusters and in macrophages that persist in the forebrain of WNV-recovered mice. CXCL16 is expressed on the surface of APCs, such as monocyte-derived and conventional dendritic cells, promoting immune cell chemotaxis^65, 66^. Past studies have shown that CXCL16 is important for the localization of infiltrating CD8^+^ T cells within inflamed tissues^27^. Using an established model of WNND recovery, we detected the highest levels of CXCL16 at 7 DPI, when the percentage CD8^+^CXCR6^+^ T cells are lowest in the CNS. During recovery, at 52 DPI, when almost 100% of the CD8^+^ T cells express CXCR6, the levels of CXCL16 return to mock-infected levels, which could be due the downregulation of chemokine receptors that commonly occurs after ligand binding^67^. Furthermore, we see opposite trends of CD8^+^CXCR6^+^ levels in the blood and CNS, with the highest levels of CD8^+^CXCR6^+^ T cells in the blood at 7 DPI, when CD8^+^CXCR6^+^ T cells in the forebrain are lowest, and the lowest levels of CD8^+^CXCR6^+^ T cells in the blood at 52 DPI, when CD8^+^CXCR6^+^ T cells in the forebrain are highest. Therefore, either only the infiltrated CXCR6^+^ CD8^+^ T cells from the blood persist within the CNS of WNV-recovered animals, or CXCR6 is upregulated upon CNS infiltration of CD8^+^ T cells. If the latter occurred, it would likely be due to meningeal signals^68^. Our data suggests that the former is more likely because total CD8^+^ T cell numbers decrease in the cortex and hippocampus as CXCL16 levels wane. CD8^+^ T cells are key players in virologic control and may persist as T_R_M cells in the CNS for years after recovery from neurotropic viral infections^69^ via unknown mechanisms. Published studies have demonstrated pathogen-specific T_R_M cells exhibit increased expression of CXCR6, which maintains them in the liver following malaria infection, and in the lung after influenza infection^26, 27^. Similarly, In the skin, CXCR6 expression by CD8^+^ T cells is required for optimal formation of T_R_M populations after epicutaneous infection with HSV-1 KOS^70^. Conversely, CXCR6-deficiency in the setting of infection with *Mycobacterium tuberculosis* (mTB) results in decreased bacterial burden without reduction in the number of T_R_M cells in the lungs^71^. These studies suggest that CXCR6 expression on CD8^+^ T cells has a distinct role dependent upon the type of infection. After infection with WNV, *Cxcr6^-/-^* mice exhibit similar frequency of virus-specific CD8^+^ T cells as WT mice in the forebrain, indicating no role for CXCR6 in T cell recruitment at peak CNS viral loads. In addition, *Cxcr6^-/-^* mice did not demonstrate any differences in weight loss, survival, or viral titers, suggesting that CXCR6 is not necessary for an effective immune response after WNV encephalitis, or trafficking of CD8^+^ T cells into the CNS parenchyma. By 52 DPI, however, *Cxcr6^-/-^* mice have significantly less CD8^+^CD103^+^ T cells compared to WT. While the scRNA-seq data demonstrate high expression of CXCR6 on CD4^+^ T cells in the WNV- recovered forebrain, similar analyses revealed no differences in levels of CD4^+^ T cells between WNV- infected WT and *Cxcr6^-/-^* mice at any DPI. These data suggest CXCR6 expression by CD8^+^ T cells is required to maintain and, possibly, differentiate T_R_M cells within the CNS. Indeed, adoptive transfer of CXCR6-deficient CD8^+^ T cells into *Cd8^-/-^* mice, followed by WNV infection, led to essentially no CD8^+^CD103^+^ T cells in the forebrain at 52 DPI. On the contrary, *Cd8^-/-^* mice that received WT CD8^+^ T cells exhibited high levels of CD8^+^CD103^+^ T cells at 52 DPI, 100% of which were CXCR6^+^. Thus, CXCR6 expression on CD8^+^ T cells is necessary for their maintenance as T_R_M cells within the CNS after WNV encephalitis.

We previously showed that CD8^+^ T cell-derived IFNγ signaling in microglia underlies spatial learning and cognitive deficits during recovery from WNV, and mice deficient in CD8^+^ T cells do not exhibit synapse loss during WNV infection^10^. Here, we found that 100% of all CD8^+^IFNγ^+^ T cells in WNV-recovered animals WT mice are also CXCR6^+^. Given that IFNγ produced from CD8^+^ T cells is responsible for persistent microglial activation, it is not surprising that *Cxcr6^-/-^* mice, which do not maintain CD8^+^ T cells, have decreased microglial activation at 52 DPI. We have also previously shown that WNV-recovered mice exhibit increased astrocyte expression of IL-1β within the hippocampus^52^. In this study we show that *Cxcr6^-/-^* mice have attenuated astrocytic production of IL-1β in the hippocampus at 52 DPI compared to WT WNV-infected mice, suggesting T_R_M cells are important for astrocyte activation. While it has been shown that CXCL16 can act on astrocytes to promote hippocampal neuroprotection against excitotoxic damage^22^, our data suggests that this effect is likely due to lack of microglial activation, as, in previous studies, we found that this associated with lack of IL-1β expression by astrocytes^10^.

Finally, as CD8^+^ T cells are necessary for viral clearance, but their persistence in the CNS can lead to chronic microglial activation and long-term cognitive sequelae^10, 31^, therapies that restrict T_R_M generation without disrupting CD8^+^ T cell antiviral functions may be valuable. To test this, we administered neutralizing anti-murine CXCL16 antibodies (Ab) during peak infection. We found anti-CXCL16 Ab, but not administration of isotype control IgG, decreased the percentages of CD8^+^CD103^+^ T cells in the forebrain of WNV-infected mice, without affecting the percentages of CD8^+^ T cells. This suggests that the CXCL16/CXCR6 axis could be an important target to reduce T_R_M cell differentiation without affecting CD8^+^ trafficking into the CNS.

## Conclusions

In summary, our studies identified unique, activated microglia clusters that persist in the CNS after recovery from viral encephalitis, with transcriptional signatures that suggest antigen-presenting capability or inflammatory function similar to those seen in aging and neurodegenerative diseases. We further identified CXCL16/CXCR6 factors as necessary for the maintenance of T_R_M cells in forebrain of mice during recovery from WNV infection, contributing to microglia and astrocyte activation, and synapse elimination in the CA3 region of the hippocampus. As initial activation of glial cells and infiltration of antiviral CD8^+^ T cells into the CNS are critical for viral clearance, identification of mechanisms responsible for persistent neuroinflammation during recovery are essential for the development of successful therapeutics to prevent post-infectious cognitive sequelae without affecting viral clearance. The CXCL16/CXCR6 chemokine signaling pathway may also be an important target for other neurological and neurodegenerative diseases that are associated with neuroinflammation in the CNS.

## Methods

### Mice

8- to 10-week old male and female mice were used for all experiments. C57BL/6J mice and *Cxcr6^-/-^* and *Cd8^-/-^* mice were obtained from Jackson Laboratories. Transgenic mice were back- crossed more than ten generations to C57BL/6 mice at Jackson Laboratories. All experiments followed guidelines approved by the Washington University School of Medicine Animals Safety Committee (protocol no. 20180120).

### Mouse model of WNV infection

M. Diamond at Washington University in St. Louis provided WNV-NS5-E218A strain utilized for intracranial infections. WNV-NS5-E218A contains a single point mutation in the gene encoding 2′-O-methyltransferase. Deeply anesthetized mice were intracranially administered 1×10^4^ plaque-forming units (p.f.u.) of WNV-NS5-E218A into the third ventricle of the brain with a guided 29-guage needle. Viruses were diluted in 10µl of 0.5% fetal bovine serum in Hank’s balanced salt solution (HBBS, Gibco). Mock-infected mice were intracranially injected with viral diluent.

### scRNA-seq

Mice (*n*=2 per group, 4 mice pooled per *n*) were deeply anesthetized and perfused intracardially with ice-cold dPBS (Gibco). Forebrain tissue were aseptically dissected, minced and enzymatic digested in HBSS (Gibco) containing collagenase D (Sigma, 50mg/ml), TLCK trypsin inhibitor (Sigma, 100μg/ml), DNase I (Sigma, 100U/μl), HEPES 7.2 (Gibco, 1M), for 1hr at 37°C while shaking. The tissue was pushed through a 70μm strainer and spun down at 500*g* for 10min. To remove myelin debris, cells were resuspended in 37% Percoll and spun at 1,200g for 30 min. To minimize cell loss and aggregation, cells were resuspended at 100 cells/μL in PBS containing 0.04% (w/v) BSA. ∼17,500 cells were partitioned into nanoliter-scale Gel Bead-In-EMulsions (GEMs) to achieve single cell resolution for a maximum of 10,000 individual cells/sample. Polyadenylated mRNA from an individual cell was tagged with a unique 16 basepair 10x barcode and 10 basepair Unique Molecular Identifier utilizing the v2 Chromium Single Cell 3′ Library Kit and Chromium instrument (10x Genomics). Full length cDNA was amplified to generate sufficient mass for library construction. cDNA amplicon size (∼400 basepair) for the library was optimized using enzymatic fragmentation and size selection. The final library was sequence-ready and contained four unique sample indexes. The concentration of the 10x single cell library was determined via qPCR (Kapa Biosystems). The libraries were normalized, pooled, and sequenced on the HiSeq4000 platform (Illumina). Five single cell libraries were sequenced across an entire HiSeq4000 flow cell targeting ∼45,000 reads per cell.

### Alignment and barcode assignment

The Cell Ranger Single-Cell Software Suite was utilized to perform sample demultiplexing, barcode processing, and single-cell 3′ counting. Cellranger mkfastq was used to demultiplex raw base call files from the HiSeq4000 sequencer into sample-specific fastq files. Files were demultiplexed with 98%^+^ perfect barcode match, and 74%^+^ q30 reads. To align reads to the mm10 mouse genemoe, fastq files for each sample were processed with cellranger counts. Samples were subsampled to have equal numbers of confidently mapped reads per cell. 4,250 total cells were counted after filtration and cluster removal.

### Preprocessing analysis with Seurat package

The Seurat package was used to analyze the processed scRNA data. Filtered genes by barcode expression matrices from Cell Ranger were used as analysis inputs. The merge function was used to pool samples. Cells with a mitochondrial fraction > 5% or less than 150 gene reads were filtered out. Expression measurements for each cell were normalized by total expression and then scaled to 10,000, and log-transformed. Two sources of unwanted variation (total UMI counts and the fraction of mitochondrial reads) were removed withthe “vars.to.regress” option in the ScaleData function.

### Dimensionality reduction and clustering

The Seurat FindVariableGenes function was used to identify the most variable genes, which were then used for dimension reduction using PCA analysis. T-SNE (t-distributed Stochastic Neighbor Embedding) plots were used to visualize cells. For clustering, we used FindClusters that utilizes shared nearest neighbors (SNN) and modularity optimization. This is based off the clustering algorithm with 20 PCA components and a clustering resolution of 0.3.

### Identification of cluster-specific genes and marker-based classification

**The Seurat** FindAllMarkers function was used with the default Wilcoxon rank sum test for single cell gene expression to identify marker genes. For each cluster, only genes that were expressed in more than 35% of cells with at least 0.25-fold difference were considered. Major CNS cell-type identities for each cluster was assigned by cross-referencing these genes with published gene expression datasets (ref). Differential expression tests between clusters were performed using the FindMarkers function using the Wilcoxon Rank Sum test, with significant genes identified as with adjusted p values <= 0.05.

### Microglial clusters Analysis

After assigning cell type identities to the clusters, differential expression tests were performed comparing clusters 0 (homeostatic microglia), 1 (activated micor-glia-1) and 4 (activated microglia-2)1, with the seurat findMarkers function using the default options and Wilcoxon Rank Sum test. A Venn diagram has been created to show the numbers of significantly differential expression genes with adjust p values of <= 0.05 from pairwise compari- sons of the 3 clusters. For the heat maps, differential expression tests were performed comparing cluster 0 to the combined cells of clusters 1 and 4, and between clusters 1 and 4, with the seurat findMarkers function using the default options and Wilcoxon Rank Sum test. The top 25 up-regulated and 25 down-regulated genes were selected based on false discovery rate (FDR) <=0.05.

### Pathway analysis

Generally applicable gene set enrichment (GAGE) method was used^72^ based on the log2 fold changes from the single cell differential expression analysis.

### Flow cytometry

Murine forebrain cells were isolated and stained with fluorescence-conjugated antibodies. Briefly, mice were anesthetized and perfused with ice-cold dPBS (Gibco). Forebrain tissue were dissected out, minced, and enzymatically digested at 37°C for 1hr with shaking. The digestion buffer contained collagenase D (Sigma, 50mg/ml), TLCK trypsin inhibitor (Sigma, 100μg/ml), DNase I (Sigma, 100U/μl), HEPES buffer, pH7.2 (Gibco, 1M) in HBSS (Gibco). The tissue was then pushed through a 70μm strainer and pelleted by a 500g spin cycle for 10min. Cells were resuspended in 37% Percoll and spun at 1,200g for 30min to remove myelin debris and then resuspended in FACS buffer. Cell were blocked for 5min at 4°C with TruStain fcX (BioLegend, cat. no. 101320), followed by cell surface staining for 15min at 4°C and cell fixation/permeabilization for 30min at 4°C. Cells were blocked again for 10min at room temperature followed by intracellular staining for 10min at room temperature and resuspended in 2% PFA for data acquisition. Data were collected using a Fortessa X-20 instrument and analyzed with the software FlowJo.

### Antibodies for flow cytometry

The following antibodies were used for flow cytometry: CD11b (Brilliant Violet 605, BioLegend, cat. no. 101257), CD103 (APC/Cy7, BioLegend, cat. no. 121432), CD4 (APC, BioLegend, cat. no. 100411), CD45 (PerCP/Cy5.5, BioLegend, cat. no. 103132), CD8 (Brilliant Violet 711, BioLegend, cat. no. 100748), CXCR6 (PE, BioLegend, cat. no. 151103), Foxp3 (PerCP-Cy5.5, Biosciences, cat. no 563902), GFP (AF488, BioLegend, cat. no. 338008), and I-A/I-E (MHC-II, APC/Cy7, BioLegend, cat. no. 107628), CD103 (Brilliant Violet 605, BioLegend, cat. no. 121433), CD69 (Brilliant Violet 421, BD Biosciences, cat. no. 562920), CD45 (BV737, BD Biosciences, cat. no. 748371) and LIVE/DEAD fixable blue dead cell stain kit (Invitrogen, cat. no. L23105).

### CD8^+^ T cell adoptive transfer

Spleens harvested (using aseptic technique) from naïve C57BL/6J or *Cxcr6^-/-^* mice were placed in sterile, ice-cold RPM1-1640 medium (Sigma) with 10% fetal bovine serum (FBS, Gibco). Cells were then strained into a single-cell suspension. After centrifugation (500g for 5min at 4°C), red blood cells were lysed in ACK lysis buffer (Gibco) on ice for 5min. Cells were diluted with RPMI/FBS buffer and centrifuged again. CD8^+^ T cells were isolated using negative selection CD8a^+^ T cell isolation kit (mouse, Miltenyi, cat. no. 130-095-236). CD8^+^ (4×10^6^ cells) were injected intravenously (i.v.) (0.2 mL) into male *Cd8^-/-^* mice.

### Immunohistochemistry (IHC)

Mice were anesthetized and perfused with ice-cold dPBS (Gibco), followed by ice-cold 4% paraformaldehyde (PFA). Brains were post-fixed overnight in 4% PFA and then cryopreserved in 30% sucrose (three exchanges every 24hr). Prior to tissue sectioning (10 µm), samples were frozen in OCT compound (Fischer). Tissue sections were washed with PBS and permeabilized with 0.1–0.3% Triton X-100 (Sigma-Aldrich) and blocked with 5% normal goat serum (Sigma-Aldrich) at room temperature for 1hr. Slides were then incubated in primary antibody overnight at 4°C. Slides were then incubated in secondary antibodies at room temperature for 1hr, and nuclei were counterstained with 4,6-diamidino-2-phenylindole (DAPI; Invitrogen). ProLong Gold Antifade Mountant (Thermo Fisher) was applied on slides prior to coverslips. A Zeiss LSM 880 confocal laser scanning microscope and software from Zeiss were used to acquire images. Immunofluorescent signals were quantified using the software ImageJ.

### RNAscope *in situ* hybridization (ISH/IHC)

Mice were anesthetized and perfused with ice-cold dPBS (Gibco) and post-fixed overnight in 4% PFA. Brains were then cryopreserved in 30% sucrose (three exchanges every 24hr) and embedded in OCT compound (Fischer). 10-µm sagittal tissue sections were prepared. RNAscope 2.5 HD Assay-Red (ACD, cat. No. 322360) was performed as per the manufacturer’s instructions. A probe against CXCL16 mRNA (ACD) was used. For ISH/IHC experiments, IHC procedures, as previously described, followed ISH probe detection. Images were acquired using a Zeiss LSM 880 confocal laser scanning microscope and processed using software from Zeiss. Immunofluorescent signals were quantified using the software ImageJ.

### Antibodies for immunohistochemistry

The following primary antibodies were used for IHC analyses: CD3 (1:200, BD Sciences, cat. no. 556970), GFAP (1:250; Thermo, cat. no. 13–0300, clone 2.2B10), GFP (1:200, abcam, cat. no. ab13970), IBA1 (1:200, Synaptic Systems, cat. no. 234 006), IL-1β (1:100; R&D, cat. no. AF-401), synaptophysin (1:250; Synaptic Systems, cat. no. 101004, polyclonal), CD68 (1:200; Bio-Rad, cat no. MCA1957, monoclonal), and C1q (1:200; Abcam, cat. No. ab182451, monoclonal). Secondary antibodies conjugated to Alexa-488 (Invitrogen, cat. no. A21206) or Alexa-555 (Invitrogen, cat. no. A21435) were used at a 1:200 dilution.

### Measurement of viral burden

Mice were infected with WNV and euthanized at specific days post-infection, as indicated. For tissue collection, mice were deeply anesthetized, and the brain was removed and micro-dissected. All tissues collected were weighed, and then homogenized with zirconia beads in a MagNA Lyser instrument (Roche Life Science) in 500µl of PBS, and stored at –80°C until virus titration. Thawed samples were clarified by centrifugation (2,000g at 4°C for 10min), and then diluted serially before infection of BHK21 cells. Plaque assays were overlaid with low-melting point agarose, fixed 4 days later with 10% formaldehyde, and stained with crystal violet. Viral burden was expressed on a log10 scale as p.f.u. per gram of tissue.

### Anti-CXCL16 treatment

Monoclonal Rat IgG_2A_ CXCL16 neutralizing antibody (R&D Systems, Clone # 142417, cat. no. MAB503) or monoclonal Rat IgG_2A_ isotype control (R&D Systems, Clone # 54447 cat. no. MAB006) was administered via retro-orbital injection at a dose of 25µg for 4 consecutive days on days 7, 8, 9, and 10 post infection.

### CXCL16 ELISA measurement

Tissues were collected in complete extraction buffer on ice and homogenized with an electric homogenizer. Complete extraction buffer was made of 100 mM Tris, pH7.4 (Gibco), 150mM NaCl (Gibco), 1mM EGTA (Gibco), 1mM EDTA (Gibco), 1% Triton X-100 (Sigma-Aldrich), 0.5% Sodium deoxycholate (Thermo Scientific) and protease and phosphatase inhibitor (78440, Thermo Scientific). Samples were centrifuged for 20min at 13,000rpm at 4°C. The level of CXCL16 was detected using commercial ELISA kit (DY503, R&D Systems, Inc., MN, USA), according to the manufacturer’s guidelines. The OD values at wavelength of 450 nm and 540 nm were measured by a microplate reader. Readings at 540 nm were subtracted from those of readings at 450 nm for wavelength correction. Concentrations of CXCL16 were calculated on the basis of a standard curve.

### Statistical Analysis

Prism 7.0 (GraphPad Software) was used to perform statistical analyses. All data were analyzed using an unpaired Student’s *t*-test, one-way or two-way ANOVA and correct for multiple comparisons as indicated in the corresponding figure legends. A *P* value of ≤ 0.05 was considered significant.

## Acknowledgements

We acknowledge the members of the Klein lab for helpful discussions and technical support. The authors would also like to thank W. Beatty at the Molecular Microbiology Imaging facility at Washington University School of Medicine for her imaging expertise, and Q. Li for critical reading of the manuscript. Experimental schematics were created using BioRender.com.

## Declarations

### Author Contributions

S.F.R., A.L.S., and R.S.K. designed experiments. A.L.S., S.F.R., S.A., M.K., V.A.D., and A.S. performed experiments. S.F.R., A.L.S., W.Y., J.A.M., S.P., and R.S.K analyzed the data. R.S.K., A.L.S. and S.F.R. wrote the paper. J.M and M.A. provided experiment guidance and edited the paper.

### Data and materials availability

The data from this study are tabulated in the main paper and supplementary materials. All reagents are available from R.S.K. under a material transfer agreement with Washington University. Upon publication, scRNA-seq data will be deposited in the Gene Expression Omnibus.

### Consent for publication

All authors consented to publication of this manuscript.

## Supplemental Figure Legends

**Supplemental Figure 1.**
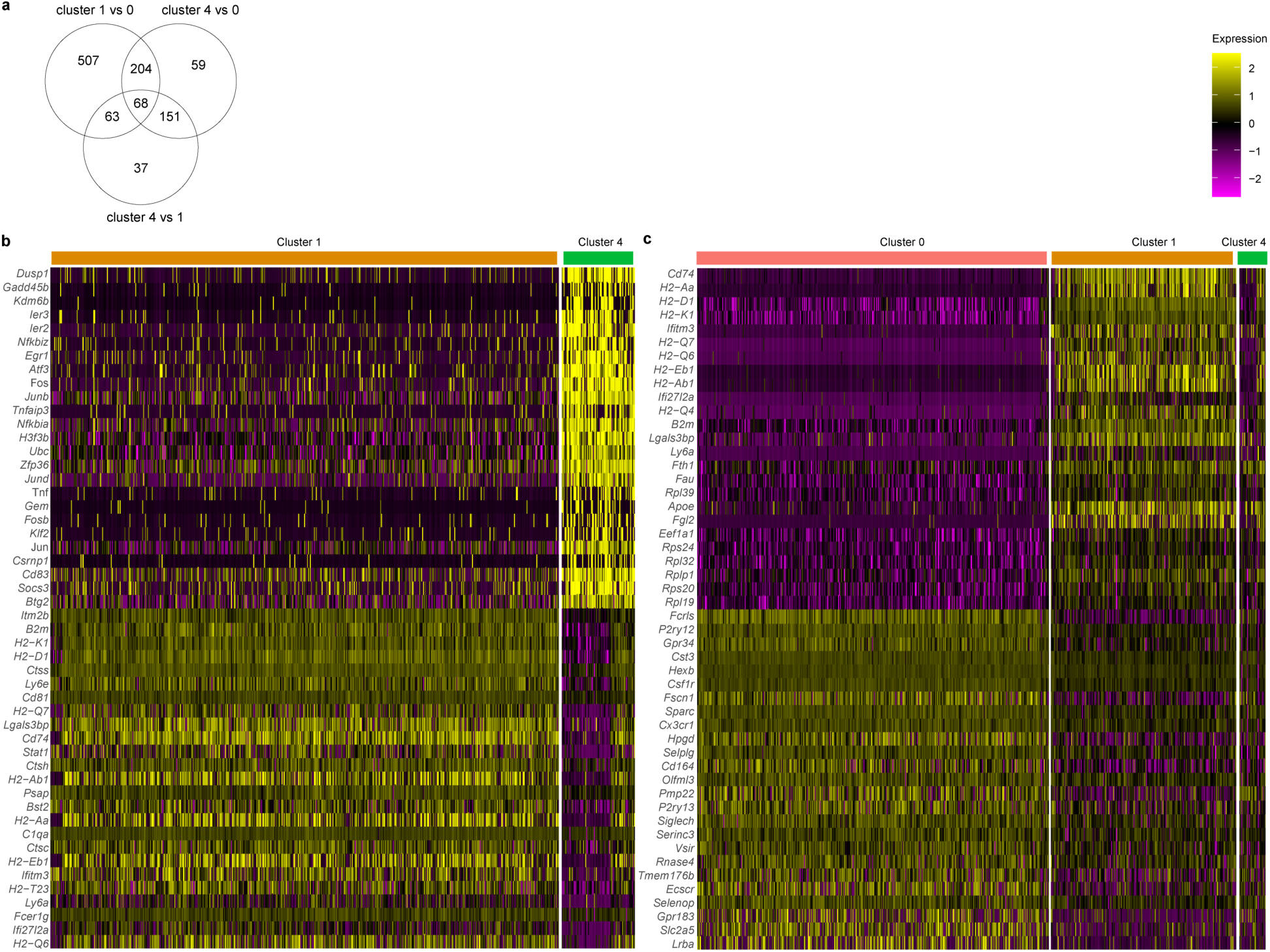
Comparative analysis of microglia clusters. **a.** Venn diagram depicting differential expression tests comparing cluster 1 to 0, 0 to 4, and 4 to 1. **b, c.** Heat maps depicting differential expression tests performed between clusters 1 and 4 (b) and cluster 0 to clusters 1 and 4 (c). False discovery rate (FDR) <=0.05.

**Supplemental Figure 2.**
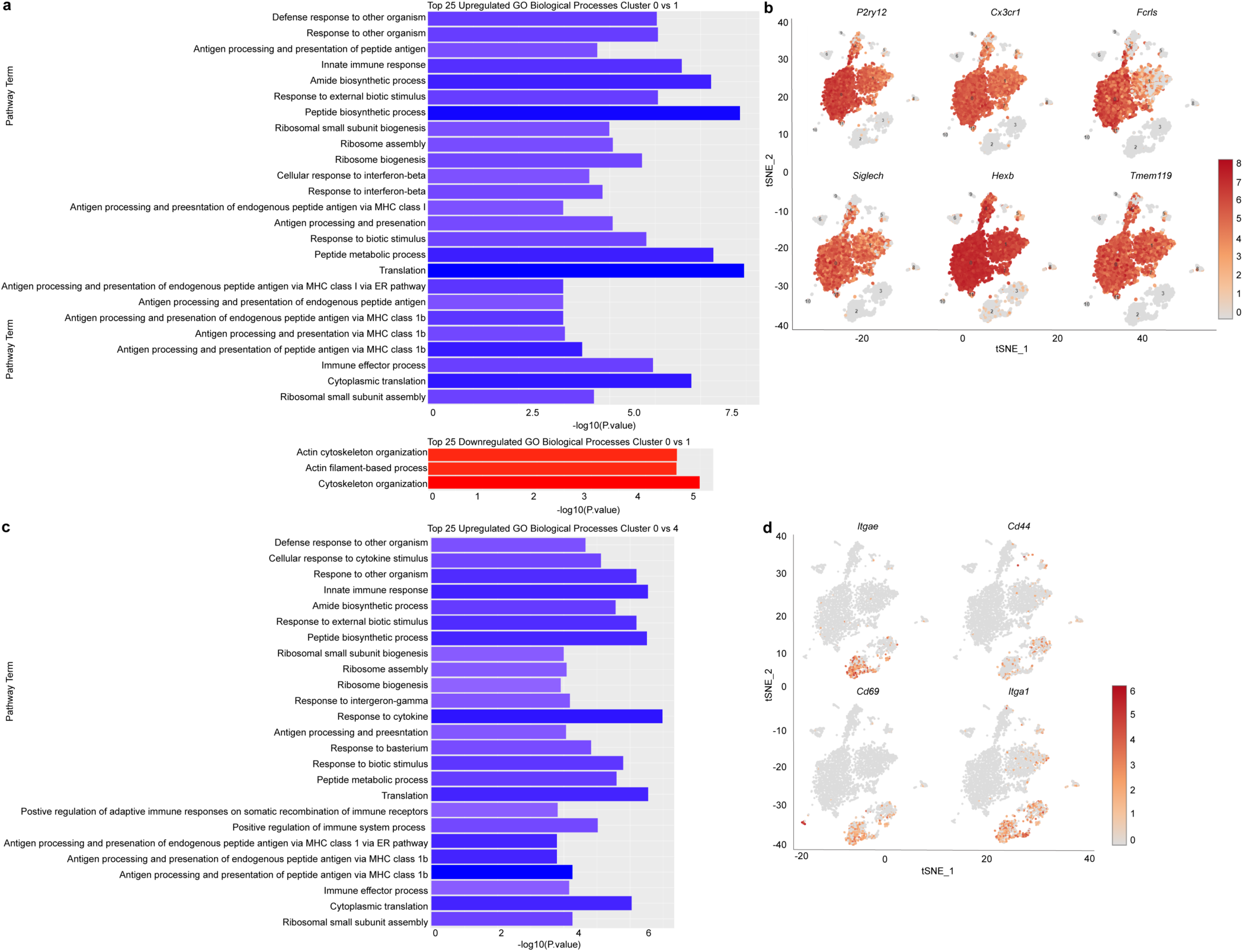
Pathway analysis of microglia clusters and tSNEs of microglia and T_R_M markers. **a, c.** GO biological processes pathway analysis using generally applicable gene set enrichment (GAGE) method^72^ based on the log2 fold changes from the single cell differential expression analysis. Pathway lists were generated with genes upregulated in Cluster 1 compared to Cluster 0 (top panel) or and downregulated in Cluster 1 compared to Cluster 0 (bottom panel) (a), or genes upregulated in cluster 4 compared to cluster 0 (c). **b.** tSNE plots depicting the relative expression of microglial core genes, *P2ry12, Cx3cr1, Fcls, Siglech, Hexb,* and *Tmem119*. **d.** tSNE plots depicting the relative expression of T_R_M genes, *Itgae, Cd44, Cd69,* and *Itga1*. Color key indicates the expression levels.

**Supplemental Figure 3.**
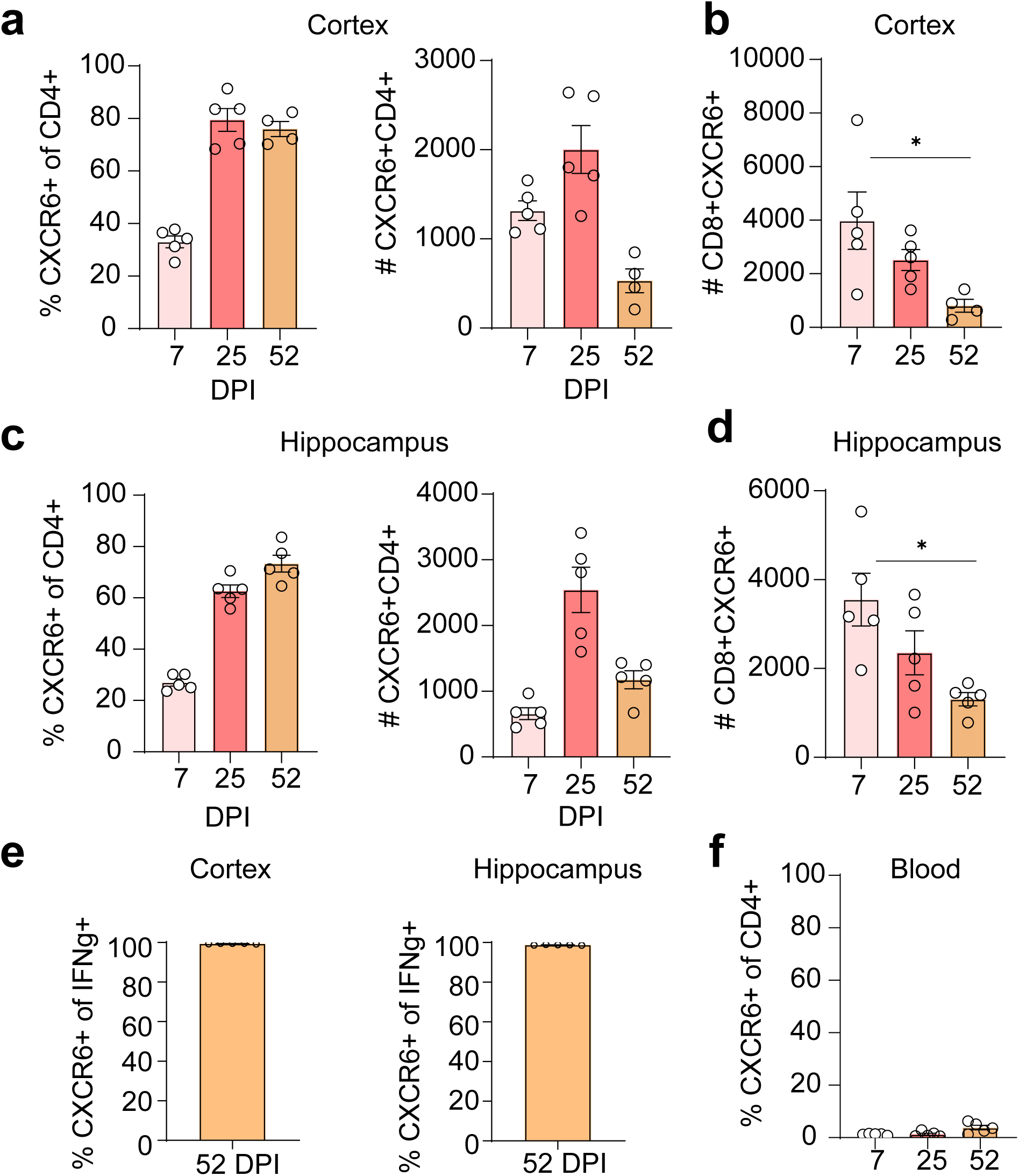
Characterization of CXCR6^+^CD4^+^ and CD8^+^ T cells during WNV infection. **a, c.** Flow cytometric analysis of the percent and number of CD4^+^ T cells that are CXCR6^+^ in the cortex (a) or hippocampus (c) of WNV-infected WT mice at 7, 25 and 52 DPI. **b,d.** Flow cytometric analysis of the number of CD8^+^ T cells that are CXCR6^+^ in the cortex (b) or hippocampus (d) of WNV-infected WT mice at 7, 25 and 52 DPI. **e.** Flow cytometric analysis of the percent of CD8^+^ T cells that are IFNg^+^ in the cortex and hippocampus of WNV-infected WT mice at 52 DPI**. f.** Flow cytometric analysis of the percent of CD4^+^ T cells that are CXCR6^+^ in the blood of WNV-infected mice at 7, 25 and 52 DPI. Data represent the mean±s.e.m. and were analyzed by unpaired Student’s *t*-test.

**Supplemental Figure 4.**
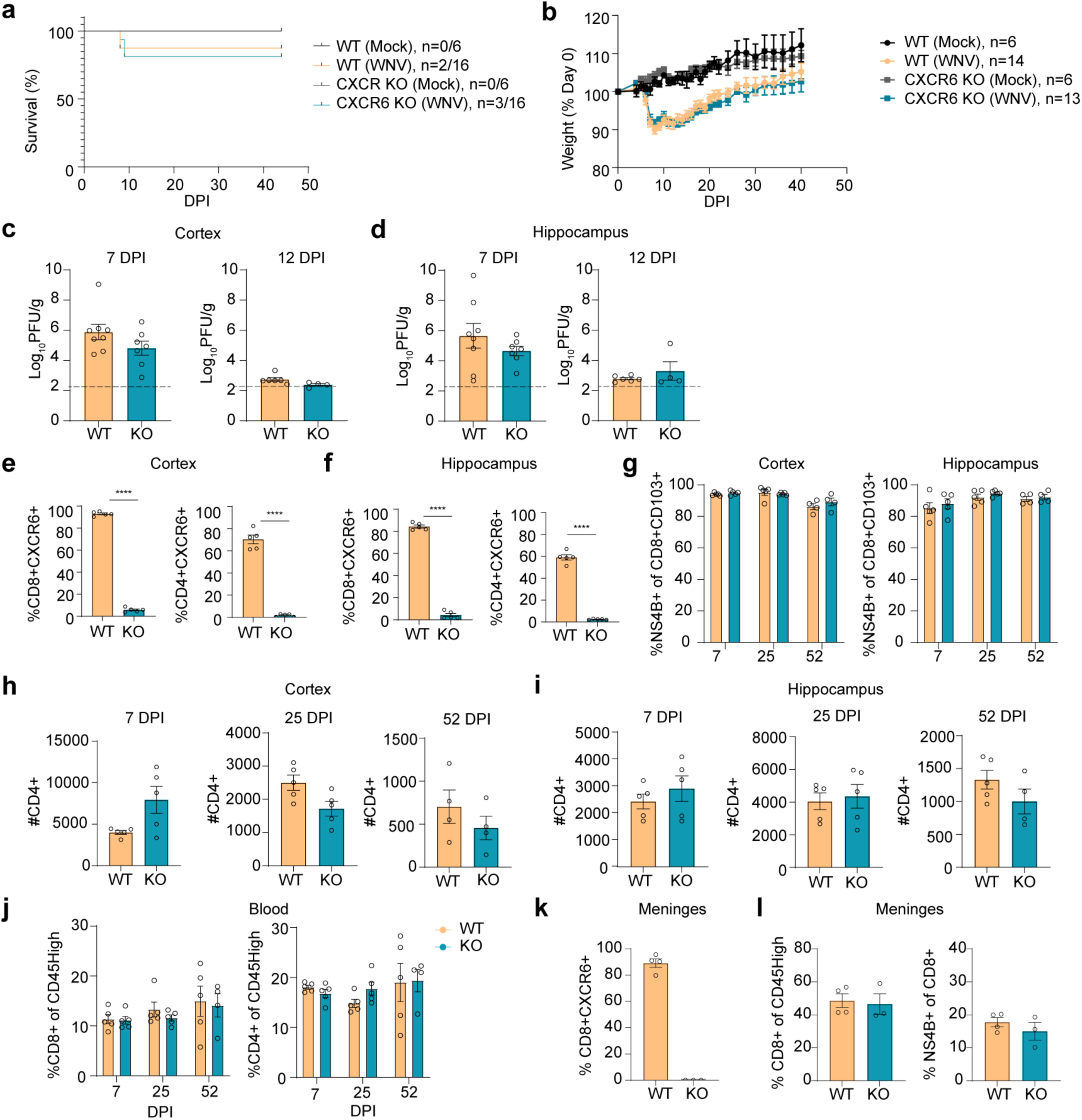
Characterization of *Cxcr6^-/-^* mice. **a.** Survival curve of WNV-infected *Cxcr6^-/-^* and WT mice. **b.** Weight loss course in WNV-infected Cx*cr6^-/-^* and WT mice. **c, d.** Viral loads measured by plaque assay in the cortex (c) and hippocampus (d) of WNV-infected *Cxcr6^-/-^* and WT mice at 7 or 12 DPI. **e, f.** Flow cytometric analysis of the percent of CD8^+^CXCR6^+^ and percent of CD4^+^CXCR6^+^ T cells in the cortex (e) and hippocampus (f) of WT and *Cxcr6^-/-^* animals at 25 DPI. **g.** Flow cytometric analysis of the percent of CD8^+^CD103^+^ T cells that are NS4B^+^ in the cortex and hippocampus of WNV-infected WT and *Cxcr6^-/-^* at 7, 25, and 52 DPI. **h,i.** Flow cytometric analysis of the total number of CD4^+^ T cells in the cortex (h) and hippocampus (i) of WT and *Cxcr6^-/-^* animals at 7, 25 and 52 DPI. **j.** Flow cytometric analysis of the percent of CD45^high^ cells that are CD8^+^ or CD4^+^ in the blood of WNV-infected Cx*cr6^-/-^* and WT mice at 7, 25 and 52 DPI. **k.** Flow cytometric analysis of the percent of CD8^+^ T cells that are CXCR6^+^ in the meninges of WNV-infected WT and Cx*cr6^-/-^* mice at 35 DPI. **l.** Flow cytometric analysis of the percent of CD45^high^ cells that are CD8^+^ followed by the percent of CD8^+^ cells that NS4B^+^ in the meninges of WT and *Cxcr6^-/-^* animals at 35 DPI. Data represent the mean±s.e.m. and were analyzed by unpaired Student’s *t*-test. \*\*\*\**P* < 0.0001.

**Supplemental Figure 5.**
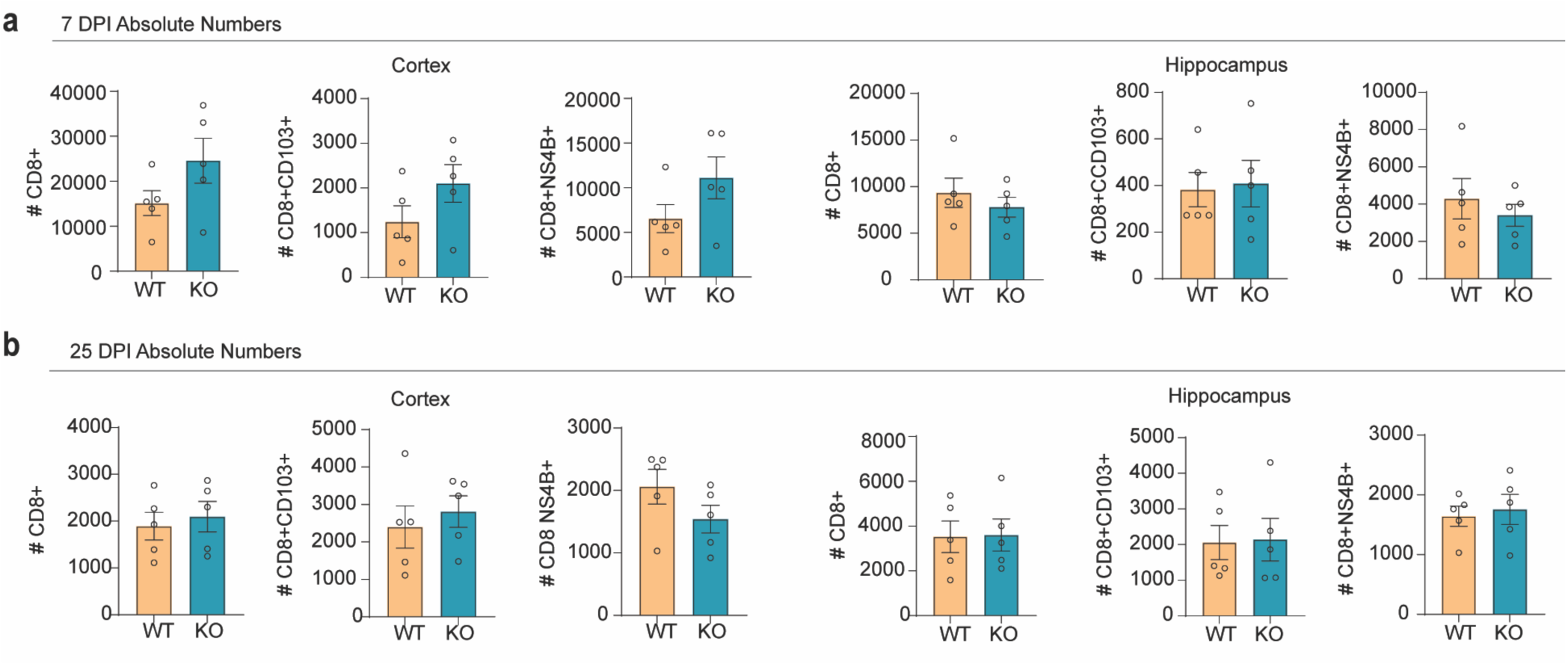
Analysis of T cell numbers in CXCR6-deficient animals. **a,b.** Flow cytometric analysis of total numbers of CD8^+^, CD8^+^CD103^+^, and CD8^+^NS4B^+^ cells in the cortex and hippocampus of WT and *Cxcr6^-/-^* mice at 7 (a) or 25 (b) DPI. Data represent the mean±s.e.m. and were analyzed by unpaired

**Supplemental Figure 6.**
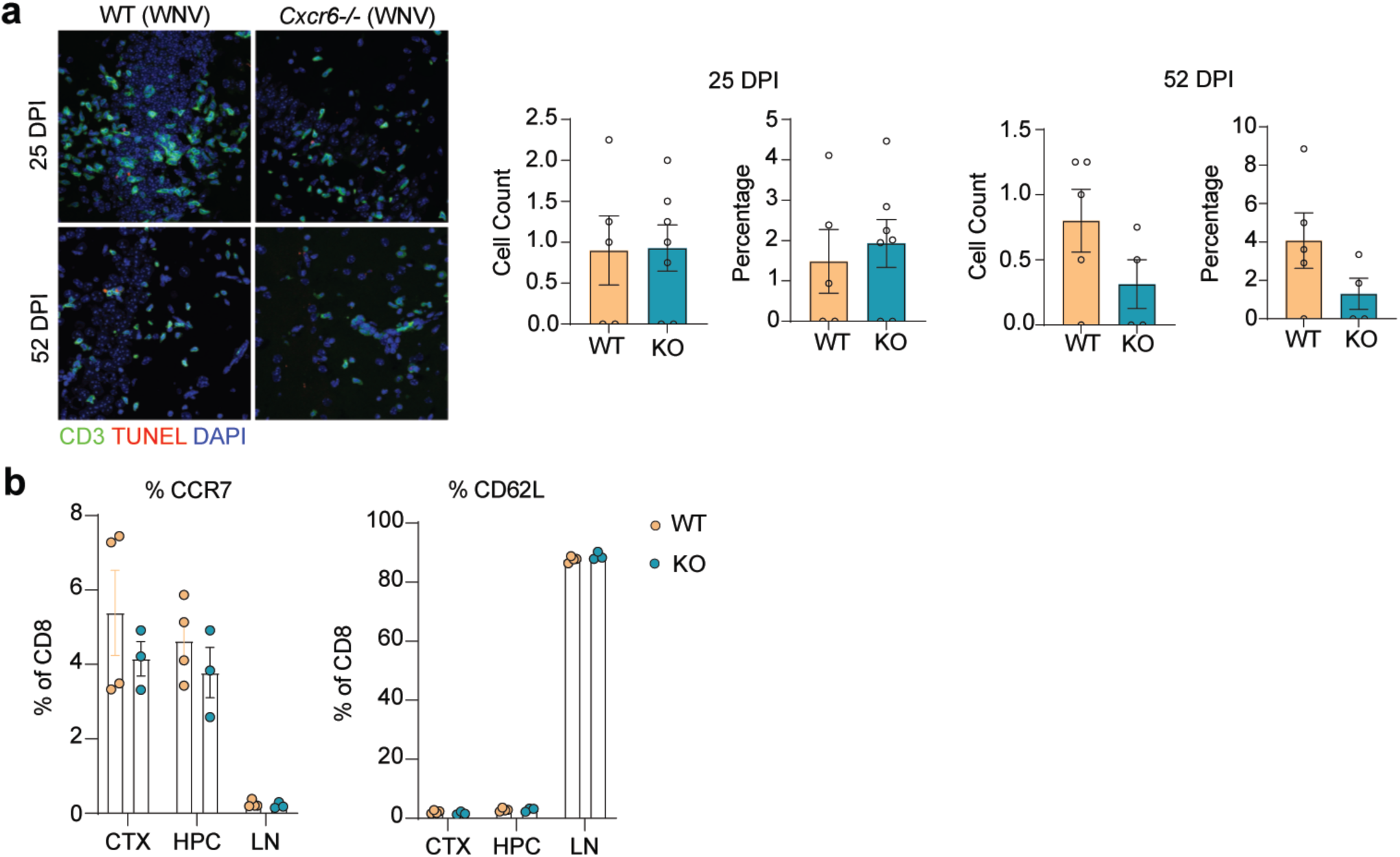
Loss of CD8^+^ T cells in the CNS is not due to T cell apoptosis, egression to the lymph node, or accumulation in the meninges in *Cxcr6^-/-^* animals. **a.** Representative immunostaining and quantification of TUNEL (red), CD3 (red) and DAPI (blue) in CA3 region of the hippocampus of WNV-infected WT or *Cxcr6^-/-^* mice at 25 and 52 DPI. Cell count quantified by counting of CD3^+^TUNEL^+^ cells per image and percentage quantified by number of CD3^+^TUNEL^+^ cells normalized to total CD3^+^ cells. **b.** Flow cytometric analysis of the percentage of CD8^+^ T cells that are CCR7^+^ or CD62L^+^ in the cortex, hippocampus, and lymph nodes at 35 DPI. Data represent the mean±s.e.m. and were analyzed by unpaired Student’s t-testStudent’s *t*-test.

**Supplemental Figure 7.**
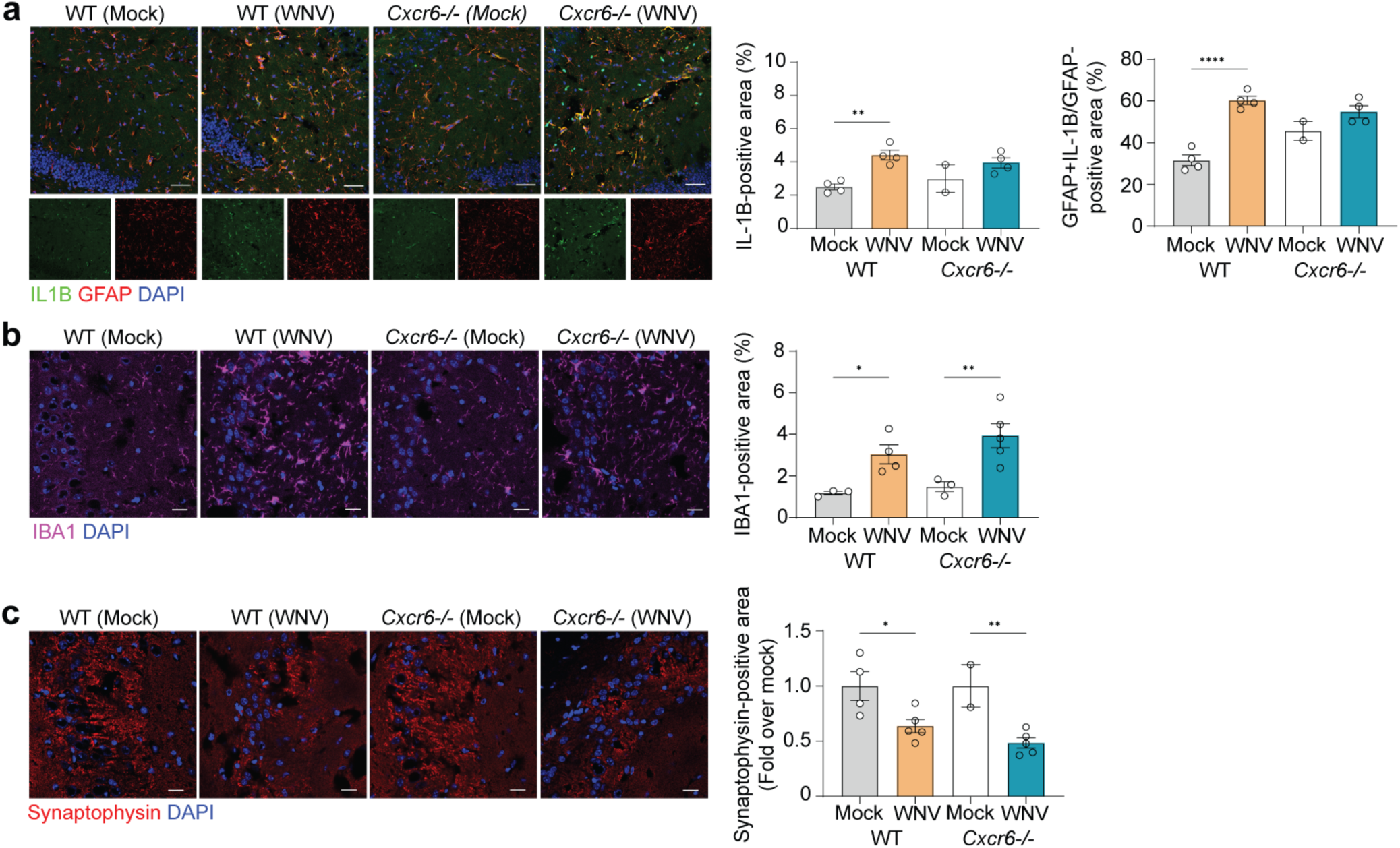
Gliosis persists in the hippocampus of WNV-infected WT and *Cxcr6^-/-^* mice at 25 DPI. **a.** Representative immunostaining at 25 DPI of IL-1β (green), GFAP (red) and DAPI (blue) in CA3 region of the hippocampus of mock or WNV-infected WT or *Cxcr6^-/-^* mice, followed by quantification of percent GFAP^+^IL-1β^+^ area, normalized to the total GFAP^+^ area. **b.** Representative immunostaining at 25 DPI of IBA1 in mock- or WNV-infected WT or *Cxcr6^-/-^* mice, showing staining for IBA1 (magenta) and DAPI (blue), followed by quantification of percent IBA1^+^ area in the hippocampus. **c.** Representative immunostaining and quantification of synapses in the CA3 region of the hippocampus in mock- or WNV-infected WT or *Cxcr6^-/-^* animals at 25 DPI showing staining for synaptophysin (red) and DAPI (blue). Synaptophysin quantified by percent positive area. Scale bars, 50 μm (a) or 20 μm (b,c). Data represent the mean±s.e.m. and were analyzed by two-way ANOVA and corrected for multiple comparisons. *P<0.05, **P<0.005, \*\*\*\**P* < 0.0001.

**Supplemental Figure 8.**
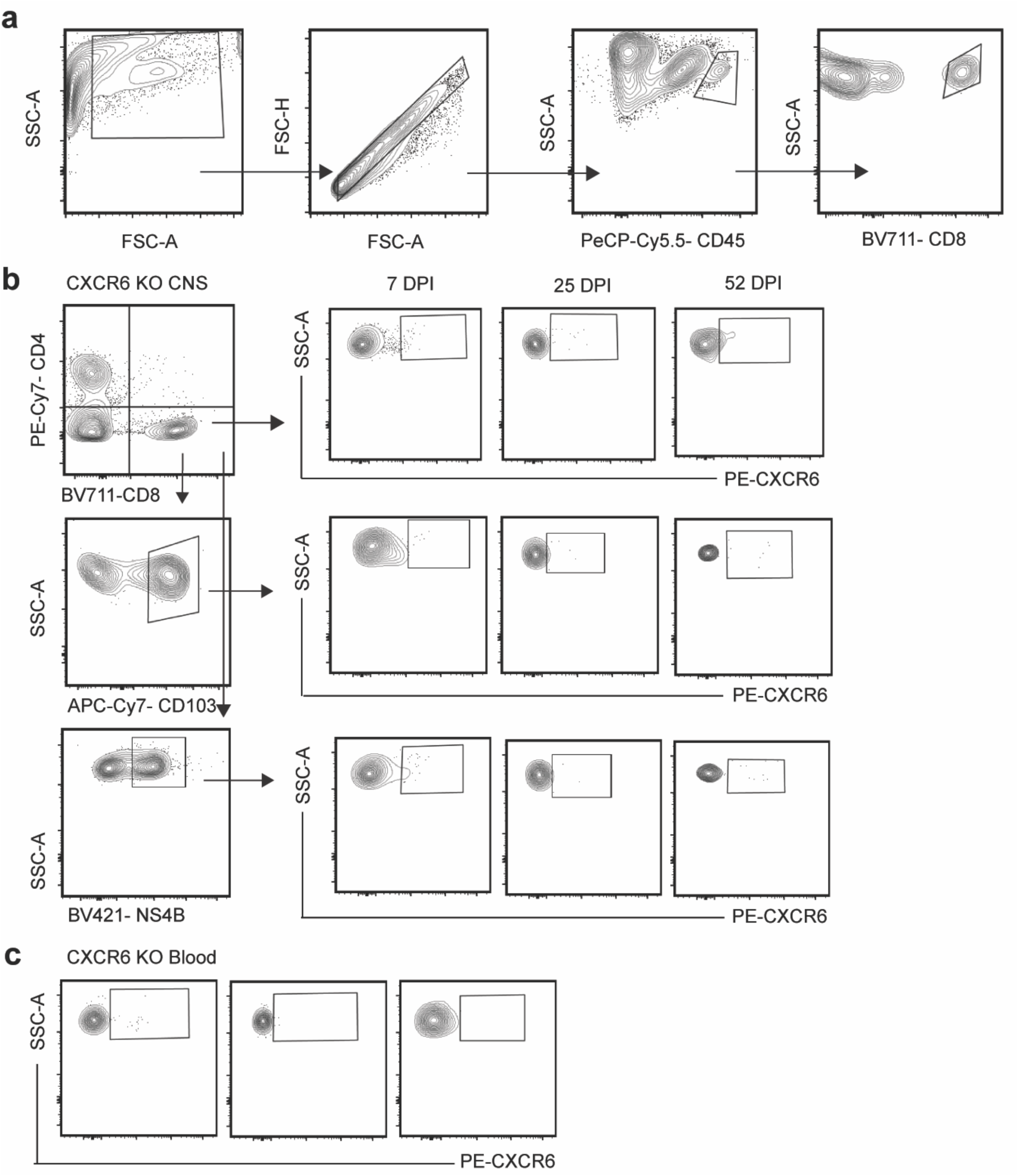
Representative gating strategies. **a.** Flow gating strategy for CD8^+^ T cells. **b,c.** Flow gating strategy to determine CXCR6^+^ cells based off gating for *Cxcr6^-/-^* mice in CNS (a) or blood (c).

---

**Supplemental Table 1.** Genes used to define cluster identity.

| Cluster |  | Gene |
| --- | --- | --- |
| 0 | Resting microglia | P2ry12, Tmem119, Siglech |
| 1 | Activated microglia -1 | Cd74, Apoe, Ccl12 |
| 2 | CD8+ T cells | Lck, Cd8a, Itgae |
| 3 | CD4+ T cells | Cd247, Cd4, Ctla4 |
| 4 | Activated microglia -2 | H2-Aa,Cst7, Ifitm3 |
| 5 | Macrophages | Cybb, Ifitm2, Mt3 |
| 6 | Astrocytes | Mt3, Fxyd1, Igtbp2 |
| 7 | IEG+ microglia | Atf3, Fos, Egr1 |
| 8 | B cells | Igkc, Mzb1, Cd79a |
| 9 | Oligodendrocytes | Alas2, Bpgm |
| 10 | Endothelial cells | Igtbp7, Car4, Ece1 |
| 11 | Microglia motility genes | Numb, Foxn3, Macf1 |

**Supplemental Table 2.**
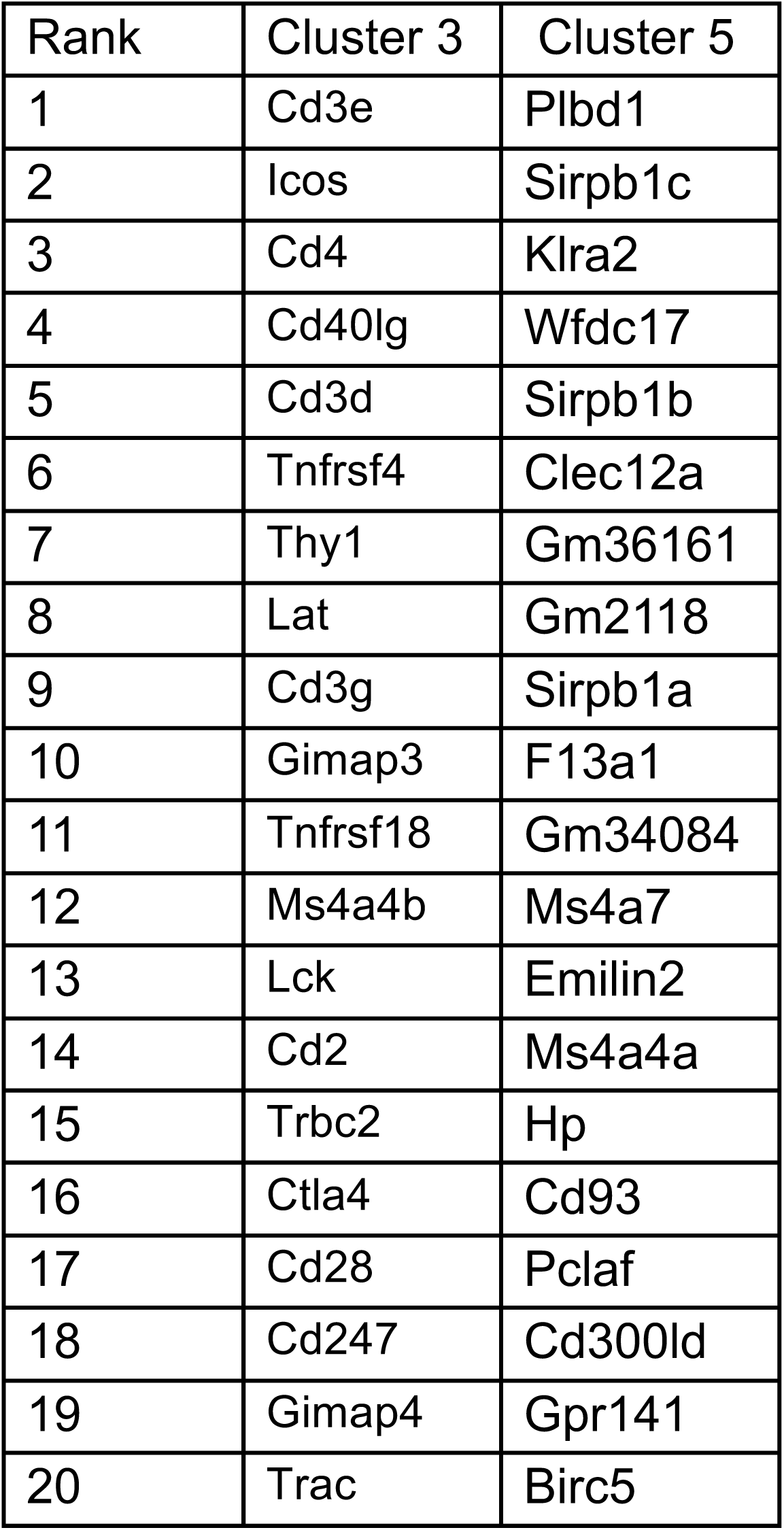
Top 20 defining genes for clusters 3 and 5.

| Rank | Cluster 3 | Cluster 5 |
| --- | --- | --- |
| 1 | Cd3e | Plbd1 |
| 2 | Icos | Sirpb1c |
| 3 | Cd4 | Klra2 |
| 4 | Cd40lg | Wfdc17 |
| 5 | Cd3d | Sirpb1b |
| 6 | Tnfrsf4 | Clec12a |
| 7 | Thy1 | Gm36161 |
| 8 | Lat | Gm2118 |
| 9 | Cd3g | Sirpb1a |
| 10 | Gimap3 | F13a1 |
| 11 | Tnfrsf18 | Gm34084 |
| 12 | Ms4a4b | Ms4a7 |
| 13 | Lck | Emilin2 |
| 14 | Cd2 | Ms4a4a |
| 15 | Trbc2 | Hp |
| 16 | Ctla4 | Cd93 |
| 17 | Cd28 | Pclaf |
| 18 | Cd247 | Cd300ld |
| 19 | Gimap4 | Gpr141 |
| 20 | Trac | Birc5 |

